# Role of neutrophil extracellular traps in regulation of lung cancer invasion and metastasis: Structural Insights from a Computational Model

**DOI:** 10.1101/2020.08.17.253575

**Authors:** Junho Lee, Donggu Lee, Sean Lawler, Yangjin Kim

**Affiliations:** Department of Mathematics, Konkuk University, Seoul, Republic of Korea; Mathematical Biosciences Institute, Ohio State University, Columbus, Ohio, United States of America; Department of neurosurgery, Brigham and Women’s Hospital & Harvard Medical School, Boston, Massachusetts, United States of America

## Abstract

Lung cancer is one of the leading causes of cancer-related deaths worldwide and is characterized by hijacking immune system for active growth and aggressive metastasis. Neutrophils, which in their original form should establish immune activities to the tumor as a first line of defense, are undermined by tumor cells to promote tumor invasion in several ways. In this study, we investigate the mutual interactions between the tumor cells and the neutrophils that facilitate tumor invasion by developing a mathematical model that involves taxis-reaction-diffusion equations for the critical components in the interaction. These include the densities of tumor and neutrophils, and the concentrations of signaling molecules and structure such as neutrophil extracellular traps (NETs). We apply the mathematical model to a Boyden invasion assay used in the experiments to demonstrate that the tumor-associated neutrophils can enhance tumor cell invasion by secreting the neutrophil elastase. We show that the model can both reproduce the major experimental observation on NET-mediated cancer invasion and make several important predictions to guide future experiments with the goal of the development of new anti-tumor strategies. Moreover, using this model, we investigate the fundamental mechanism of NET-mediated invasion of cancer cells and the impact of internal and external heterogeneity on the migration patterning of tumour cells and their response to different treatment schedules.

**Author summary:** When cancer patients are diagnosed with tumours at a primary site, the cancer cells are often found in the blood or already metastasized to the secondary sites in other organs. These metastatic cancer cells are more resistant to major anti-cancer therapies, and lead to the low survival probability. Until recently, the role of neutrophils, specifically tumor-associated neutrophils as a member of complex tumor microenvironment, has been ignored for a long time due to technical difficulties in tumor biology but these neutrophils are emerging as an important player in regulation of tumor invasion and metastasis. The mutual interaction between a tumor and neutrophils from bone marrow or in blood induces the critical transition of the naive form, called the N1 type, to the more aggressive phenotype, called the N2 TANs, which then promotes tumor invasion. In this article, we investigate how stimulated neutrophils with different N1 and N2 landscapes shape the metastatic potential of the lung cancers. Our simulation framework is designed for boyden invasion chamber in experiments and based on a mathematical model that describes how tumor cells interact with neutrophils and N2 TANs can promote tumor cell invasion. We demonstrate that the efficacy of anti-tumor (anti-invasion) drugs depend on this critical communication and N1 → N2 landscapes of stimulated neutrophils.

## Introduction

Lung cancer is still the leading cause of cancer-associated deaths worldwide, with 1.8 million deaths in 2018 [1, 2]. Various cell types such as immune cells, fibroblasts, and endothelial cells in a tumor microenvironment (TME) interact with tumor cells via the cytokines and growth factors. Tumor-associated neutrophils (TANs) are of particular interest because experimental studies showed that they can contribute to the tumor growth, critical invasion, epithelial-mesenchymal transition (EMT), and metastasis of cancer cells [3, 4]. Until recently, neutrophils have been considered as merely a bystander in the TME and metastasis [5–7] but they are emerging as an important player due to consistent and continuous evidences of their tumor-promoting roles [3]. The high neutrophil to lymphocyte ratio (NLR) is considered to be an important indicator of poor prognosis of various cancers [3] including lung cancers [8, 9]. It was shown that lung cancer cells can elaborate CXC chemokines attracting neutrophils to TME [10] and neutrophil invasion is highly correlated with poor clinical outcomes [11, 12]. Neutrophils are the most abundant leukocytes in blood that play important roles in our innate immune system by killing harmful microorganisms via three mechanisms [13]: (i) phagocytosis (engulfing and digestion of bacteria or fungi), (ii) degranulation of cytotoxic enzymes, (iii) neutrophil extracellular traps (NETs), which consist of DNA meshes with cytotoxic enzymes and trap microorganisms in the extracellular space. Therefore, the classical form of neutrophils, called N1 TAN, can induce lysis of tumor cells [14–16] through many mechanisms including reactive oxygen species (ROS)-mediated hypochlorous acid [17, 18] and kill tumor cells via TNF-*α* [19]. Furthermore, TANs induced from interferons (IFNs) can secrete tumour necrosis factor-related apoptosis-inducing ligand (TRAIL) which then binds to a receptor of tumor cells and passes it to downstream pathways for tumor cell apoptosis [20]. However, tumor-supporting TANs are recruited from tumor cells at a distant TME and infiltrate tumor tissue, leading to tumor growth, invasion, metastasis [21–25] and ultimately, poor clinical outcomes in many cancers [26]. Metastatic cancer cells were able to induce neutrophils to form metastasis-promoting NETs without involving infection processes [27].

While transforming growth factor (TGF-*β*) is known to facilitate tumor growth, invasion, and metastasis of many types of cancers [28, 29] including breast [30–32] and lung [28] cancers, the tumor-secreted TGF-*β* was shown to transform N1 TANs (tumor-suppressive phenotype) to N2 TANs (tumor-promoting phenotype) [33–35]. The reverse action of this transformation can be mediated by type I IFN [19, 33], which can inhibit tumor growth [36] through regulation of TYK2, JAK1, STAT family, and their downstream pathways [37].

Excess activity of neutrophil elastase (NE or ELANE) has been shown to cause tissue damage and to harmful remodeling process in lung diseases such as pneumonia, acute lung injury, and cancer [38]. Importantly, NE was shown to infiltrate the TME [39] and promote tumor growth in lung cancers through the PIK3 signaling pathways [10]. Neutrophil can also promote the tumor cell invasion by degrading the extracellular matrix (ECM) and basal membrane with NE [40] and matrix metallopeptidase (MMP) [17, 41, 42]. NE was also suggested to induce EMT and metastasis of cancer cells [4, 41, 42]. For example, NE inhibitors were able to inhibit tumor growth and metastasis [40, 43, 44]. TANs were also suggested to drive angiogenesis in malignancy by secreting MMPs, which subsequently promote VEGF secretion [17]. It was shown that neutrophils can promote the tumor cell invasion in the transwell assay [27, 45] and *in vivo* experiments [27, 46, 46].

The detailed mechanism of tumor invasion and metastasis via communication with TANs is still poorly understood (Fig 1). It would be difficult to build a comprehensive mathematical model of the tumor invasion and metastasis that incorporates all the biochemical and mechanical processes in Fig 1. As a beginning step, we focus on the neutrophil-mediated invasion of tumor cells, for which there are experimental data.

**Fig 1.**
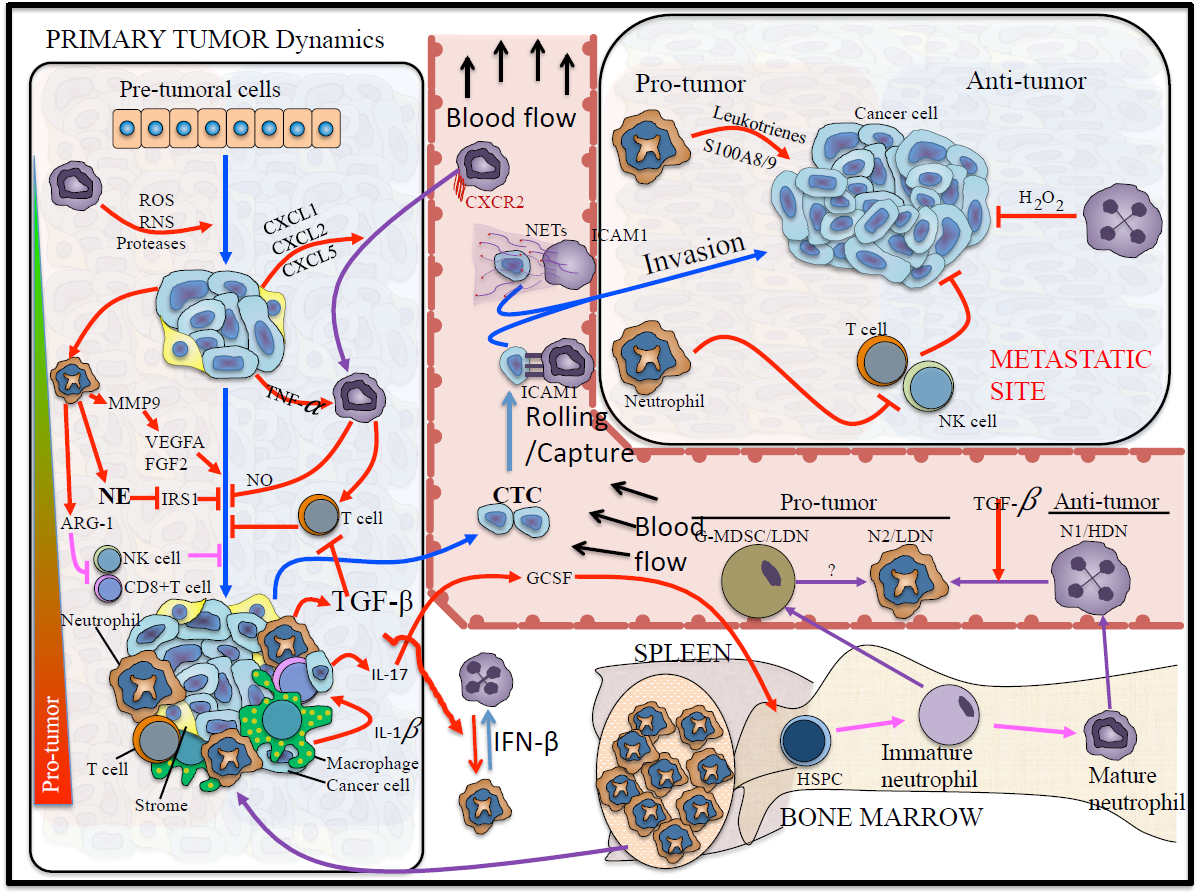
Schematic of tumor-TAN interactions for tumor growth and metastasis [47–53].

Here, we develop a mathematical model based on taxis-reaction-diffusion equations that govern cell-cell signaling and chemotactic cell movement. Our goal is to understand the biochemical factors that are important in regulating the chemotactic movement of tumor cells from the upper chamber to the lower well of the Boyden chamber assay shown in Fig 3A. We show that the mathematical model can replicate the major components of experimental findings and we test several anti-invasion intervention strategies with predictions.

**Fig 2.**
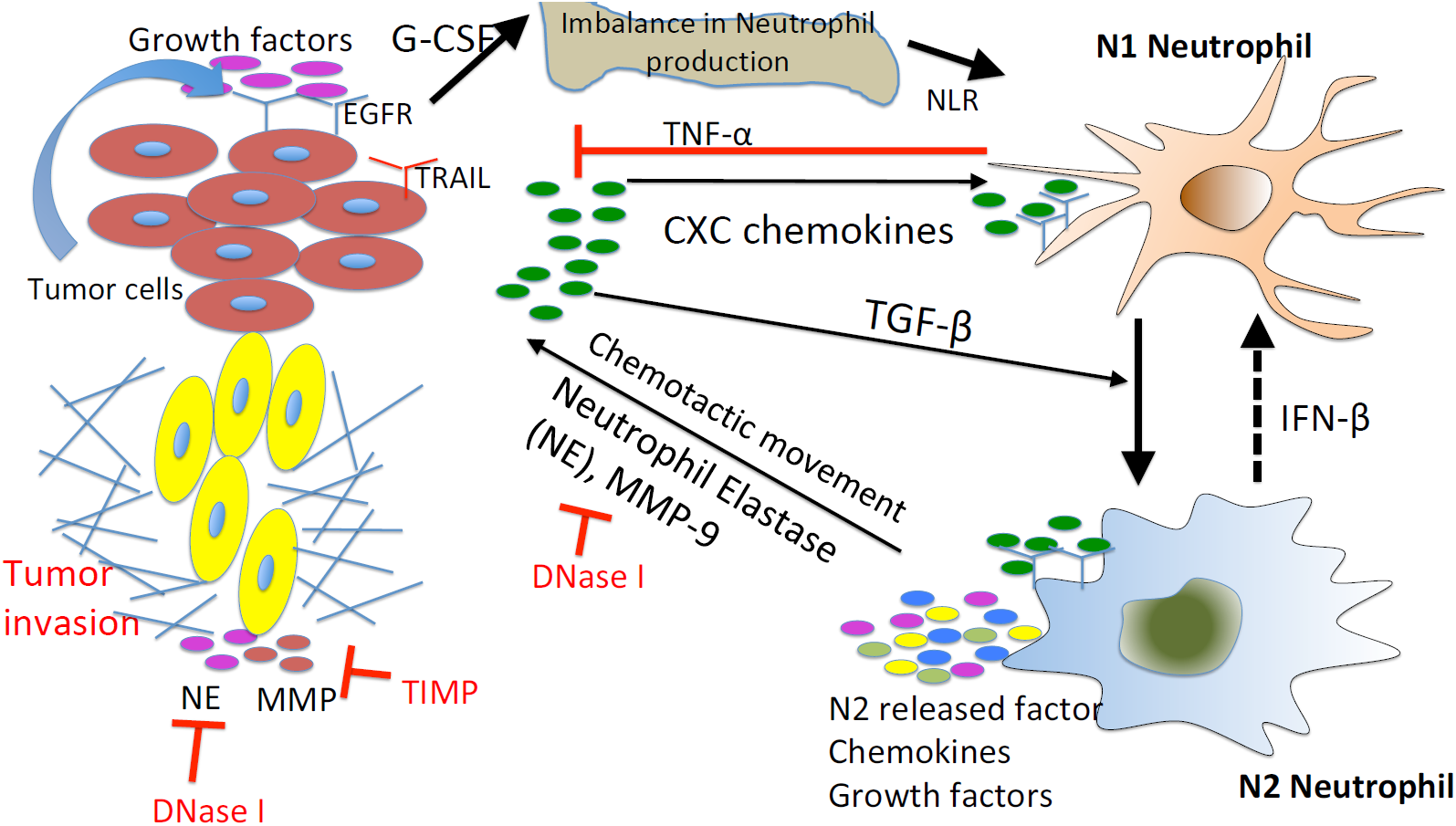
Interaction of the TGF-*β*, IFN-*β*, and NE-pathways in the control of tumor cell invasion. In homeostasis of normal tissue, these pathways are balanced so as to control growth, but in lung cancer increased secretion of TGF-*β* by tumor cells induces the N1 → N2 transition of the neutrophils and stimulates their secretion of NE and other growth factors. This disrupts the homeostasis and stimulates aggressive tumor invasion.

**Fig 3.**
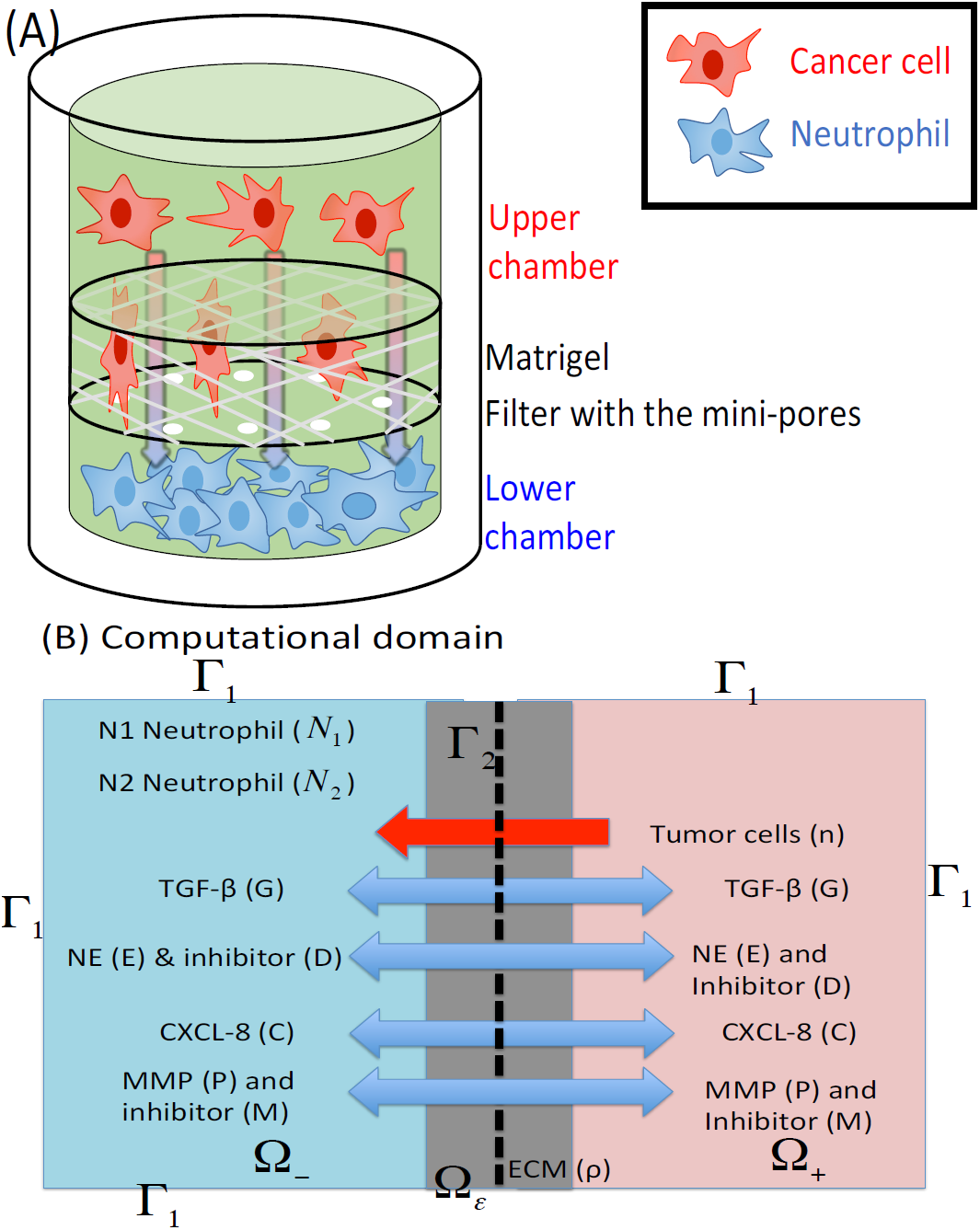
Schematics of an Invasion Assay System : (A) Boyden transwell invasion assay. Tumor cells were suspended in the upper chamber, while neutrophils or medium alone (control) were placed in the lower chamber. Semipermeable inserts coated with Matrigel ECM were inserted in the filter. In response to NE secreted by N2 neutrophils in the lower chamber, tumor cells degrade the heavy ECM proteolytically and invade the lower chamber. The number of neutrophils on the lower surface of the permeable insert was counted after 22 *h* in the absence and presence of neutrophils in the lower chamber. (B) TGF-*β* (*G*), NE (*E*), NE inhibitor (*D*), CXCL-8 (*C*), MMP (*P*), TIMP (*M*) and tumor cells (*n*) can cross the semi-permeable membrane, but neither type of neutrophils (*N*_1_, *N*_2_) can cross it. Initially, the tumor cells reside in the upper chamber (domain Ω_+_) while neutrophils are placed in the lower chamber (domain Ω_−_). An ECM layer surrounds the filter, semi-permeable membrane.

## Materials and Methods

We developed a mathematical model of tumor cell invasion in *in vitro* experiments, a critical step in metastasis [27, 54, 55], based on mutual interactions between tumor cells and TANs (Fig 2).

We denote by Ω the 3-dimensional domain

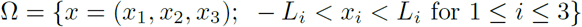

and set

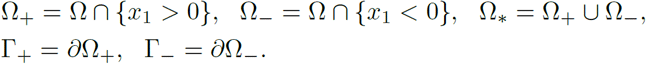

The semi-permeable membrane occupies the planar region

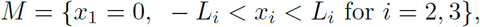

and the ECM occupies a 3-dimensional region

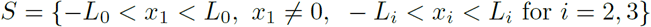

where 0 *< L*_0_ *< L*_1_. We denote by *I*_*A*_ the characteristic function of a set *A* :

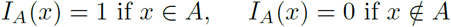

The geometry of the experimental setup of the Boyden invasion chamber was shown in Fig 3A. In the typical transwell migration assay, neutrophils isolated from the bone marrow are plated in the lower chamber, and tumor cells were added on top of Matrigel-coated insert in the upper chamber [27]. In our model, we assume that tumor cells are initially placed on the top of ECM coated area above the membrane with mini-pores in the middle, and invade the lower chamber where neutrophils (or conventional medium for control) reside. The corresponding computational domain is shown in Fig 3B.

We introduce the following variables at space **x** and time *t*:

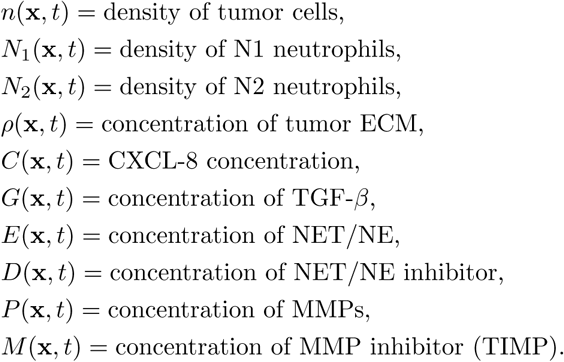

The evolution equations for these variables are developed in next sections, but in this work we focus on the Boyden invasion chamber, transwell assay, in one space dimension.

### Tumor cell density (= *n*(x, *t*))

The mass balance equation for the tumor cell density *n*(**x**, *t*) is

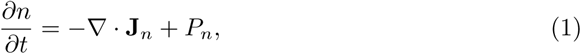

where **J**_*n*_ is the flux and *P*_*n*_ is the net production rate of cancer cells. The flux **J**_*n*_ is comprised of three components, **J**_*random*_, **J**_*chemo*_, and **J**_*hapto*_, which are the fluxes due to random motion, chemotaxis, and haptotaxis, respectively [31, 56].

We assume that the tumor ECM is homogeneous and isotropic in TME, and that the flux due to the random motility is given by

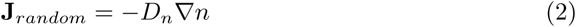

where *D*_*n*_ is the random motility constant of tumor cells.

In lung tissue, tumor cells are strongly attracted to chemotactic attractants [46] such as NE and neutrophils [27, 46] and migrate toward the up-gradient (∇*E*) of the chemo-attractant, NE, through the process called *‘chemotaxis’* [57]. The chemotactic flux is assumed to be of the form

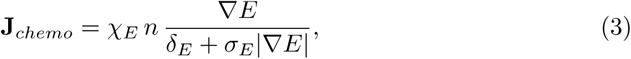

where *χ*_*E*_ is the chemotactic sensitivity, *δ*_*E*_, *σ*_*E*_ are scaling parameters, and *E* is the concentration of NE, whose dynamics will be introduced in Section below. This form reduces to the standard form of the chemotactic flux (**J**_*chemo*_ ≈ *C n* ∇*E*; *C*=constant) under small NE gradients (|∇*E*| « 1) and saturates (**J**_*chemo*_ ≈ (*χ*_*E*_*/δ*_*E*_) *n* **u**; **u** = ∇*E/*|∇*E*| is the unit vector) under large NE gradients, preventing the blow-up behaviors of solutions [31]. Other forms such as 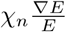[58] or 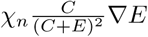 (*C*: constant) [59] have been adapted in the literature. See [58–62] for analysis of blow-up behaviors of solutions in some class of chemotaxis models.

Tumor cell invasiveness is enhanced by proteolytic degradation of the ECM via MMPs [4, 35, 63] and NEs [40, 41] that are produced by neutrophils. This results in local degradation of tumor ECM [56] and tumor cell movement in the direction of the up-gradient (∇*ρ*) of ECM via a cellular process called *haptotaxis*. This process is valid only in the ECM domain *S*, therefore, we include the characteristic function *I*_*S*_, providing the on-off switch on the ECM membrane. We represent the haptotactic flux in a similar fashion:

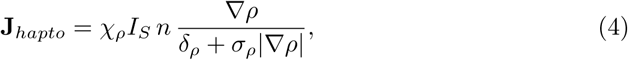

where *χ*_*ρ*_ is the haptotactic sensitivity, *δ*_*ρ*_, *σ*_*ρ*_ are scaling parameters, and *ρ* is the concentration of tumor ECM, whose dynamics will be introduced in Section below.

The net production of tumor cells is due to active NE-stimulated growth [10, 27] and cell killing by N1 TANs [3, 26, 33], which we represent as follows.

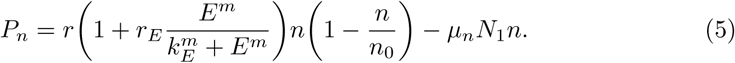

Here *r* is the proliferation rate of tumor cells in the absence of NE (*E*), *r*_*E*_ is the dimensionless parameter of NE-mediated tumor growth, *k*_*E*_ and *m* are Hill-function coefficients for activation of proliferation in the presence of NE, *n*_0_ is the carrying capacity of the tumor in a given TME, and, finally *μ*_*n*_ is the killing rate of tumor cells by N1 neutrophils (*N*_1_) whose dynamics will be described in Section below. Here, *r, r*_*E*_, *k*_*E*_, *n*_0_, *μ*_*n*_ ∈ ℝ^+^, *m* ∈ ℤ^+^.

Combining the several fluxes in Eqs (2)-(4) and growth term in Eq (5) leads to the governing equation for the tumor cell density

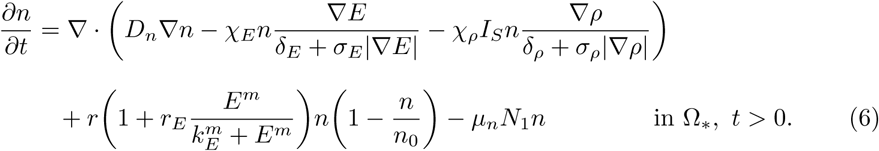

### Densities of neutrophils: N1 (= *N*_1_(x, *t*)) & N2 (= *N*_2_(x, *t*)) types

We use a similar form of reaction-diffusion-advection equations for the evolution of the densities of neutrophils, based on mass balance as in the previous section. We assume that (i) Neutrophils are chemotactic to the CXCL secreted by tumor cells [64–66], and the chemotactic flux is of the nonlinear form (3), but with different chemotactic sensitivities (*χ*_1_, *χ*_2_). Since the N2 TANs produce NE and MMPs, the movement of activated neutrophils further enhances tumor invasiveness and growth via the NE-PI3K pathway described earlier. (ii) The anti-tumorigenic (N1) neutrophils transform into the active N2 type at the rate *λ*_12_ in the presence of TGF-*β*, based on experimental evidences [3, 26, 67]. For instance, the N1 → N2 transition of TANs with protumour properties was typically observed in a TGF-*β*-rich TME and the presence of IFN-*β* or TGF-*β* inhibitor can mediate the reverse transition (N2 → N1) with anti-tumoral properties [3]. Therefore, TGF-*β* pathway inhibitors are under clinical trials since they were shown to promote the development of N1 TANs [68, 69]. (iii) N1 and N2 phenotypes proliferate at a rate, *λ*_1_ and *λ*_2_(*G*), respectively. Then, we have the following evolution equations:

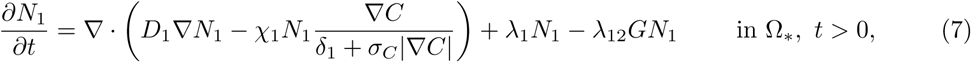

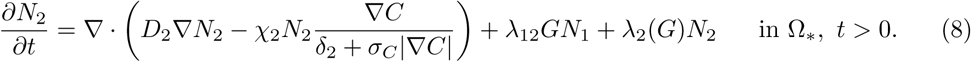

### Tumor ECM density (=*ρ*(x, *t*))

The tumor ECM provides structural foundation for efficient cell migration [70], but it also needs to be remodeled via proteolysis for tumor cell migration by microenvironmental proteases [64, 71–73]. In this work, we assume that the tumor ECM is degraded by the TAN-secreted NEs [74] and TAN-secreted MMPs [4, 35, 63, 64] as in the invasion experiments [27]. The rate of ECM change can be represented as

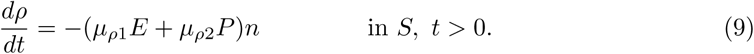

Here *μ*_*ρ*1_, *μ*_*ρ*2_ are the degradation rates by NEs and MMPs, respectively, which are secreted by N2 neutrophils. Essentially, this equation represents proteolytic degradation of tumor ECM coated on the filter when there is a significant level of tumor ECM present, as is normally the case in a TME.

### CXCL-8 concentration (= *C*(x, *t*))

Tumor cells secrete CXCL in order to recruit the immune cells such as neutrophils [64–66]. CXCLs and corresponding receptors (CXCR) such as CXCL5 and CXCR6 are important prognostic factors, alone or in a combination with the TANs, for shorter overall survival and cumulative risk of recurrence [3, 75, 76]. Thus the governing equation for CXCL8 is:

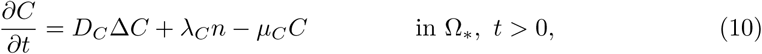

where *D*_*C*_ is the diffusion coefficient of CXCL, *λ*_*C*_ is the secretion rate of CXCL by tumor cells and *μ*_*C*_ is the decay rate of CXCL.

### TGF-*β* concentration (= *G*(x, *t*))

TGF-*β* is a polypeptide that plays a major role in regulation of many human diseases including cancers [29] due to its capacity of maintaining tissue homoeostasis and involving in most of the chronic inflammatory and wounding processes by activating from its inactive form in ECM [77]. Tumor cells is the primary source of TGF-*β* in TME [3, 48]. TGF-*β* activates proinflammatory and antitumorigenic N1 neutrophils into the aggressive N2 type, which in turn stimulates tumor cell invasion [47, 64]. Thus the governing equation for TGF-*β* is as follows.

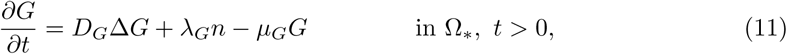

where *D*_*G*_ is the diffusion coefficient of TGF-*β, λ*_*G*_ is the secretion rate of TGF-*β* by tumor cells, and *μ*_*G*_ is the decay rate of TGF-*β*.

### Concentrations of NET/NE (= *E*(x, *t*)) and its inhibitors (= *D*(x, *t*))

NET and NE are highly associated with aggressive invasion, growth, EMT, and metastasis of cancer cells [4, 41, 42]. NE is produced by neutrophils [4, 74, 78] and used for degradation of ECM and tissue destruction [10, 41, 79]. It was also shown that NE inhibitors such as DNase I blocks this effect in growth model [10] and invasion assays [27]. In our framework, NET and the associated NEs are merged into one component. Thus, the governing equations for NET/NE and its inhibitors are

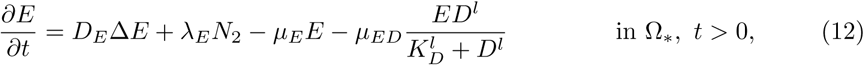

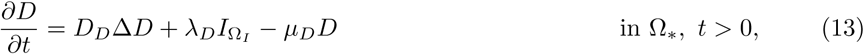

where *D*_*E*_, *D*_*D*_ are diffusion coefficients of NET/NE and its inhibitors, respectively, *λ*_*E*_ is the production rate of NET/NE from N2 neutrophils, *λ*_*D*_ is the injection rate of NET/NE inhibitors at a subdomain Ω_*I*_, *μ*_*E*_, *μ*_*D*_ are natural decay rates of NET/NE and its inhibitors, respectively, *μ*_*ED*_ is the consumption rate of NE in response to NE inhibitors with kinetic parameters *K*_*D*_, *l* (*D*_*E*_, *D*_*D*_, *λ*_*E*_, *λ*_*D*_, *μ*_*E*_, *μ*_*D*_, *μ*_*ED*_, *K*_*D*_ ∈ ℝ^+^, *l* ∈ ℤ^+^).

### MMP concentration (= *P* (x, *t*))

MMPs are highly associated with cancer cell invasion and metastasis [55, 80]. Neutrophils, not tumor cells [64, 81], were suggested to the primary source of MMPs [4, 35, 63, 64] including MMP-9 [63] in lung cancer development, showing strikingly predominant presence at the invasive fronts of metastatic cancers [63]. Thus the governing equation for MMPs is

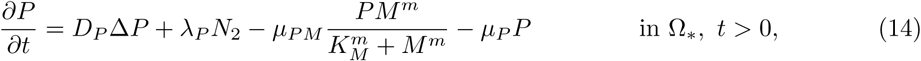

where *D*_*P*_ is the diffusion coefficient of MMPs, *λ*_*P*_ is the MMP production rate by N2 neutrophils, *μ*_*P M*_ is the degradation of MMPs by its inhibitor, TIMP, with Hill-coefficients *K*_*M*_, *m* (*K*_*M*_ ∈ ℝ^+^, *m* ∈ ℤ^+^), *μ*_*P*_ is the decay rate of MMPs. In general, *D*_*P*_ is very small (*D*_*P*_ « 1) while the half-life of MMPs is short (*μ*_*P*_ » 1) [82], leading localized activities at the moving front of invasive cells.

### TIMP concentration (= *M* (x, *t*))

TIMPs play an important role in inhibiting tumor invasion and metastasis [83] by regulating major signalling pathways in pericellular proteolysis of various ECMs and cell surface proteins [84]. In the model, TIMPs are injected for inhibition of the proteolytic activities of cancer cell invasion. Note, however, that this action can partially block cancer cell invasion since cancer cells can still execute the NE-mediated invasion. Thus, the governing equation of TIMP is

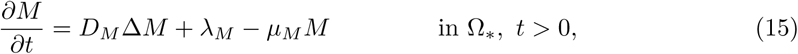

where *D*_*M*_ is the diffusion coefficient, *λ*_*M*_ is the TIMP supply rate, and *μ*_*M*_ is the decay rate of TIMP.

### Boundary conditions and initial conditions

In the following simulations we prescribe Neumann boundary conditions on the exterior boundary Γ_1_ (see Fig 3B) as follows:

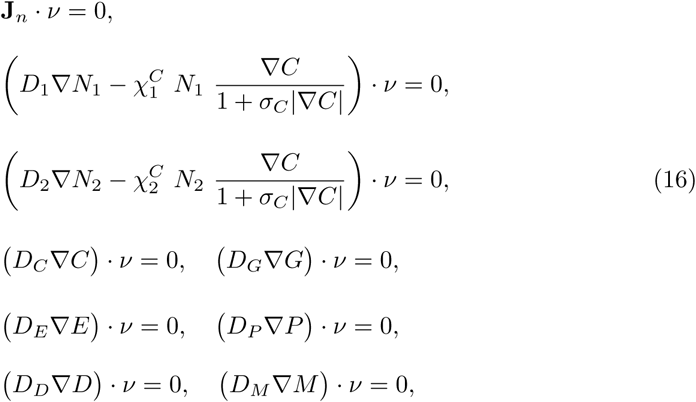

where *v* is the unit outer normal vector. The membrane is permeable to all variables (*n, N*_1_, *N*_2_, *C, G, E, D, P, M*), but not freely so. We describe the flux at membrane boundary Γ_2_ (see Fig 3B) for these variables **u** = (*n, N*_1_,*N*_2_, *C, G, E, D, P, M*) as

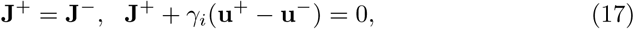

where

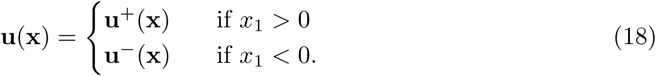

Here, the parameters *γ*_*i*_ (*γ*_*i*_ *>* 0, *i* = 1, …, 9) represent the permeability of cells (*i* = 1, 2, 3) and molecules (*i* = 4, *…*, 9). The permeability (*γ*_*i*_) is determined by the density and size of the holes on the semi-permeable membrane between upper and lower chambers as well as the size of the moving object relative to the hole size. The holes in the insert are uniformly distributed on the membrane of the Boyden invasion transwell assay [27, 45]. See [124] for the derivation of these Robin-type boundary conditions by the homogenization method. If the size of the circular holes in the membrane is increased (or decreased), the membrane becomes more (or less) permeable, and *γ*_*i*_ increases (or decreases) [30, 31]. For instance, the diameter of typical cells is in the range of 10-20*μm* while the size of growth factors and cytokines such as TGF-*β* is much smaller [125]. Furthermore, the diffusion coefficient of cells is usually much smaller than that of growth factors and cytokines [126]. While, the typical diffusion coefficient of molecules (CXCL8, EGF, and TGF-*β*) is in the range of (1.0-2.5)×10^−6^ *cm*^2^*/s* [66, 89–93], the random motility coefficient of cells is much smaller ((1.0-10.0)×10^−10^ *cm*^2^*/s*). Therefore, the parameter (*γ*_*i*_ = *γ*_*c*_ (*i* = 1, 2, 3)) of the migratory cells is smaller than the permeability parameter (*γ*_*i*_ = *γ* (*i* = 4, *…*, 9)) of the diffusible molecules due to different physical sizes. In a classical Boyden invasion chamber, a typical, invasive tumor cell in the upper chamber is not able to invade the lower chamber if the diameter of the permeable holes on the membrane is less than 0.4*μm* while molecules can diffuse throughout the domain [30]. So, we take the smaller permeability parameter for cells (*γ*_*c*_ *< γ*).

Finally, we prescribe initial conditions,

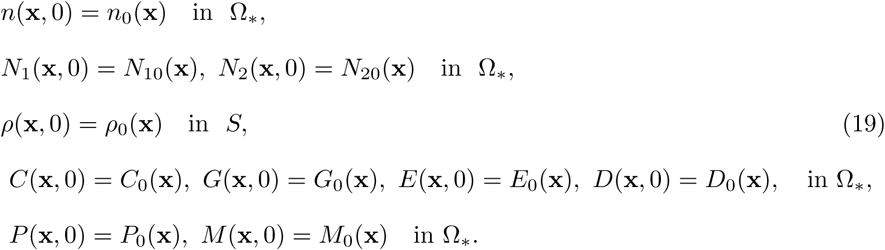

Parameters are given in Tables 1-2. Parameter estimation and nondimensionalization of the system (6)-(19) are given in S1 Appendix. This non-dimensional form of governing equations was used for the simulations. Hereafter, the computational domain is restricted to one space dimension, and the computational domain is scaled to unit length.

**Table 1.** Parameters used in the tumor model.

|  | Description | Dimensional Value | Refs. |
| --- | --- | --- | --- |
| Diffusion coefficients ($cm^2s^{-1}$) | | | |
| $D_n$ | Tumor cells | $2.5 \times 10^{-8}$ | [30, 85–88] |
| $D_1$ | N1 Neutrophil | $1.1 \times 10^{-8}$ | [89] |
| $D_2$ | N2 Neutrophil | $1.1 \times 10^{-8}$ | [89] |
| $D_C$ | CXCL8 (IL-8) chemokines | $2.5 \times 10^{-6}$ | [66, 89, 90] |
| $D_G$ | TGF-β | $1.0 \times 10^{-6}$ | [91–93] |
| $D_E$ | NET/NE | $5.0 \times 10^{-7}$ | [94–99] |
| $D_P$ | MMP | $5.0 \times 10^{-10}$ | [88, 100, 101] |
| $D_D$ | DNase I (NET/NE inhibitor) | $7.374 \times 10^{-6}$ | [102], estimated |
| $D_M$ | TIMP (MMP inhibitor) | $8.33 \times 10^{-7}$ | estimated |
| $D_A$ | TGF-β Anti-body | $8.33 \times 10^{-7}$ | estimated |
| Production rates |  |  |  |
| $r$ | Proliferation rate of tumor cells | $3.3 \times 10^{-4} s^{-1}$ | [27, 30], estimated |
| $r_E$ | NE-mediated proliferation rate of tumor cells | $7.0 \times 10^{-2}$ | estimated |
| $k_E$ | Hill type coefficient of tumor cell proliferation | $2.15 \times 10^{-9} gcm^{-3}$ | [30], estimated |
| $m$ | Hill coefficient of NE-mediated tumor proliferation | 2 | estimated |
| $n_0$ | Tumor cell carrying capacity | $2.5 \times 10^4 cells/cm^3$ | [30], estimated |
| $\lambda_1$ | Proliferation rate of N1 neutrophil | $4.38 \times 10^{-6} s^{-1}$ | [30, 103], estimated |
| $\lambda_{12}$ | Transformation rate from N1 to N2 neutrophils | $4.08 \times 10^3 cm^3g^{-1}s^{-1}$ | [30], estimated |
| $\lambda_2$ | Proliferation rate of N2 neutrophils | $2.65 \times 10^{-5} s^{-1}$ | [30, 103], estimated |
| $\lambda_C$ | Production rate of CXCL8 from tumor cells | $4.44 \times 10^{-11} s^{-1}$ | [30, 104, 105], estimated |
| $\lambda_G$ | Production rate of TGF-β from tumor cells | $4.89 \times 10^{-7} s^{-1}$ | [30, 104, 105] |
| $\lambda_E$ | Production rate of NET/NEs from Neutrophils | $2.26 \times 10^{-7} s^{-1}$ | [106] |
| $\lambda_P$ | Production rate of MMPs from Neutrophils | $2.22 \times 10^{-8} s^{-1}$ | [106] |
| $\lambda_D$ | Production rate of NET/NE inhibitor | $9.0 \times 10^{-13} gcm^{-3}s^{-1}$ | estimated |
| $\lambda_M$ | Production rate of TIMP | $1.29 \times 10^{-11} gcm^{-3}s^{-1}$ | estimated |
| $\lambda_A$ | Production rate of TGF-β anti-body | $4.78 \times 10^{-5} \mu Ms^{-1}$ | estimated |
| Degradation/Decay rates |  |  |  |
| $\mu_n$ | tumor cells degradation rate by N1 | $2.78 \times 10^{-1} cm^3g^{-1}s^{-1}$ | estimated |
| $\mu_{\rho 1}$ | ECM degradation rate by NEs | $1.02 \times 10^4 cm^3g^{-1}s^{-1}$ | estimated |
| $\mu_{\rho 2}$ | ECM degradation rate by MMPs | $3.19 \times 10^5 cm^3g^{-1}s^{-1}$ | estimated |
| $\mu_C$ | decay rate of CXCL8 | $6.42 \times 10^{-5} s^{-1}$ | [107–109] |
| $\mu_G$ | decay rate of TGF-β | $8.02 \times 10^{-6} s^{-1}$ | [30], estimated |
| $\mu_E$ | decay rate of neutrophil elastase | $8.02 \times 10^{-6} s^{-1}$ | [110] |
| $\mu_P$ | decay rate of MMP | $5.0 \times 10^{-5} s^{-1}$ | [56] |
| $\mu_D$ | decay rate of NE inhibitor | $9.627 \times 10^{-5} s^{-1}$ | [111–115] |
| $\mu_M$ | decay rate of TIMP | $4.56 \times 10^{-6} s^{-1}$ | [116] |
| $\mu_{ED}$ | degradation rate of NET/NE by its inhibitor | $(2.8 \times 10^{-4} - 2.8 \times 10^{-2}) s^{-1}$ | estimated |
| $\mu_A$ | decay rate of TGF-β anti-body | $6.42 \times 10^{-5} s^{-1}$ | [67, 117] |
| $\mu_{AG}$ | decay rate of TGF-β by anti-body | $4.8 \times 10^{-3} \mu M^{-1}s^{-1}$ | [67], estimated |
| $K_D$ | Hill coefficient of NET/NE degradation by its inhibitor | $3.2 \times 10^{-9} g/cm^3$ | [118], estimated |
| $l$ | Hill coefficient of NET/NE degradation by its inhibitor | 2 | [118], estimated |
| $\mu_{PM}$ | degradation rate of MMPs by TIMP | $(2.8 \times 10^{-5} - 2.8 \times 10^{-4}) s^{-1}$ | estimated |

**Table 2.** Parameters used in the tumor model. Continued from Table 1. : Dimensionless values were marked in ^*^.

|  | Description | Dimensional Value | Refs. |
| --- | --- | --- | --- |
| Degradation/Decay rates |  |  |  |
| $K_M$ | Hill coefficient of MMP degradation by TIMP | $4.64 \times 10^{-8} \text{ g/cm}^3$ | [118], estimated |
| $m$ | Hill coefficient of MMP degradation by TIMP | 2 | estimated |
| Movement parameters (chemotaxis, haptotaxis) |  |  |  |
| $\chi_E$ | Chemotactic sensitivity to NE | $1.117 \times 10^{-9} \text{ cm.s}^{-1}$ | [27, 30, 86, 119] |
| $\delta_E$ | Scaling parameter of NE gradient ($\vert E\vert $) | $6.46 \times 10^{-8} \text{ g/cm}^4$ | estimated |
| $\sigma_E$ | Scaling parameter of NE gradient ($\vert E\vert $) | 1.0 | estimated |
| $\chi_\rho$ | Haptotactic sensitivity | $3.5 \times 10^{-10} \text{ cm.s}^{-1}$ | [27, 120–122], estimated |
| $\delta_\rho$ | Scaling parameter of ECM gradient ($\vert \rho\vert $) | $5.0 \times 10^{-3} \text{ g/cm}^4$ | estimated |
| $\sigma_\rho$ | Scaling parameter of ECM gradient ($\vert \rho\vert $) | 1.0 | estimated |
| $\chi_1^C$ | Chemotactic sensitivity of N1 neutrophils to CXCL8 | $1.1 \times 10^{-9} \text{ cm.s}^{-1}$ | [27, 30, 123], estimated |
| $\delta_1$ | Scaling parameter of CXCL gradient ($\vert C\vert $) | $1.0 \times 10^{-11} \text{ g/cm}^4$ | estimated |
| $\chi_2^C$ | Chemotactic sensitivity of N2 neutrophils to CXCL8 | $1.1 \times 10^{-9} \text{ cm.s}^{-1}$ | [27, 30, 123], estimated |
| $\delta_2$ | Scaling parameter of CXCL gradient ($\vert C\vert $) | $1.0 \times 10^{-11} \text{ g/cm}^4$ | estimated |
| $\sigma_C$ | Scaling parameter of CXCL8 gradient ($\vert C\vert $) | 1.0* | estimated |
| Membrane parameters |  |  |  |
| $\gamma_c$ | permeability of cells (Tumor cells, N1 neutrophils, N2 neutrophils) | 0.01-785 (78.5)* | estimated |
| $\gamma$ | permeability of chemicals (CXCL8, TGF-β, NET/NE, NE inhibitor, MMP, TIMP) | 0.1-7850 (785)* | [30], estimated |
| $\mu$ | ECM width | 0.6* | estimated |

All the simulations were performed using a finite volume method (FVM; clawpack (http://www.amath.washington.edu/~claw/)) with fractional step method [127]. A nonlinear solver *nksol* was used for solving algebraic systems. The equations (6)-(19) were solved on a regular uniform grid with grid size 0.01 (*h*_*x*_=0.01). An initial time step of 0.0001 (or smaller) was used, but adaptive time stepping scheme based on the number of iterations can increase or decrease this step size.

## Results

In this section, we investigate the role of NEs in regulation of cancer cell invasion, compare the predictions of our mathematical model with experimental data, and then suggest new therapeutic strategies for blocking invasive tumor cells.

### Predictions of the mathematical model

Fig 4 shows the density profiles of all variables (*n, N*_1_, *N*_2_, *ρ, C, G, E, P*) at *t* = 0, 5, 14, 22 *h* in the absence of DNase I and TIMP when neutrophils were added in the lower chamber. In each subframe, the right (or left) half of the computational domain represents the upper (or lower) chamber in the Boyden invasion assay (Fig 3A). By degradation of the tumor ECM on the membrane of the insert, tumor cells in the upper chamber were experimentally shown to have capacity of invading the lower chamber upon stimulus of N1/N2 neutrophils in the lower chamber [27, 45]. Tumor cells in the upper chamber secrete CXCL8 (Fig 4F), which then diffuses and attracts neutrophils in the lower chamber by *chemotaxis*. On the other hand, tumor cells produce TGF-*β* (Fig 4B), which diffuses and enhances the N1 → N2 transformation of neutrophils (Fig 4C) in the lower chamber. These activated N2 TANs in the lower chamber then secrete NET/NEs (Fig 4D) and MMPs (Fig 4E) to stimulate chemotactic and haptotactic movement of tumor cells in the upper chamber. Tumor cells break down the ECM component by proteolytic activities with the NE and MMP near the membrane and invade the left chamber (Fig 4A). As they invade, they can sense higher levels of NE, and proliferate at a higher rate (See Eq (6)).

**Fig 4.**
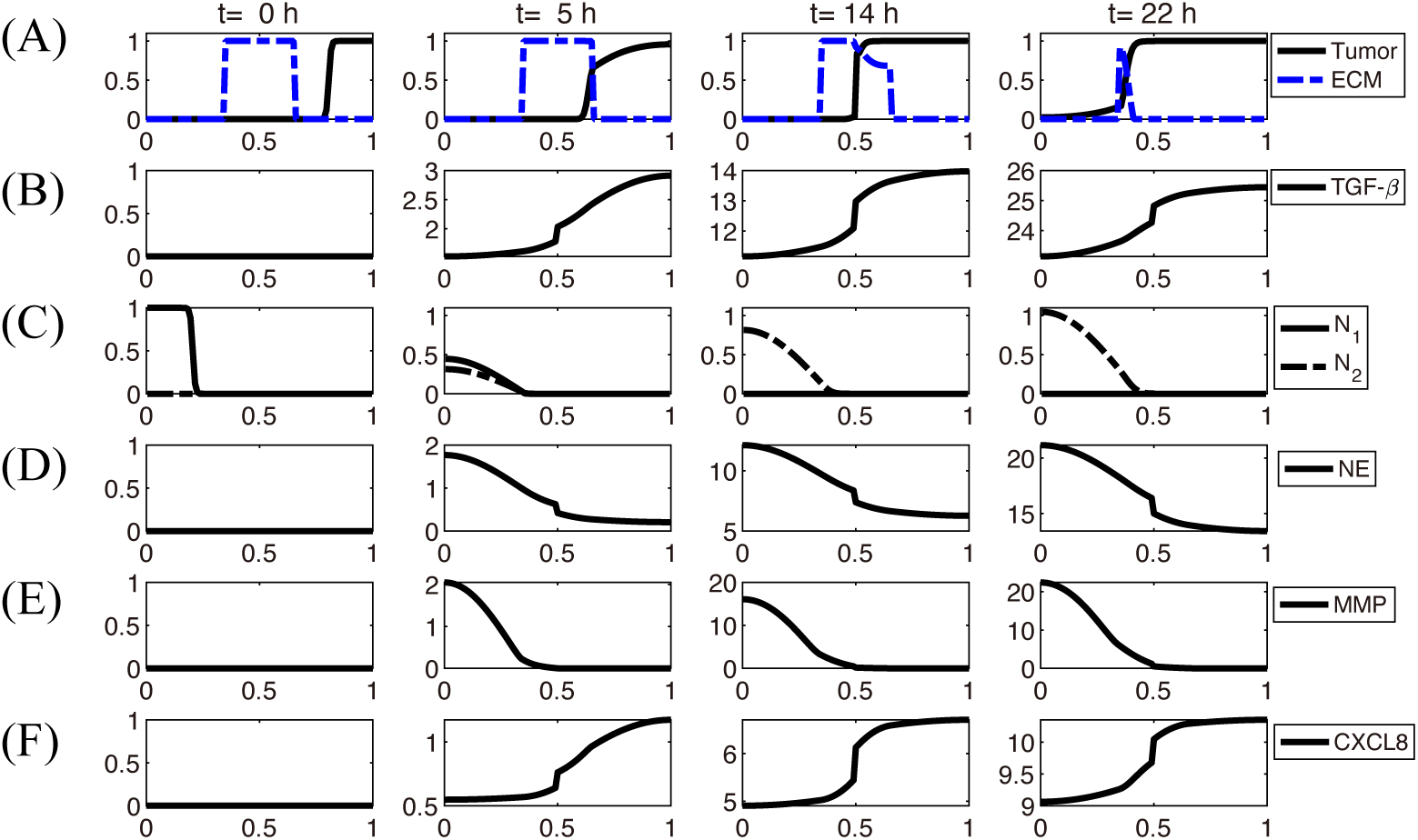
Dynamics of the system. The time evolution of the density of each variable. (A) tumor cells and ECM (B) TGF-*β* (C) N1/N2 neutrophils (D) neutrophil elastase (E) MMP (F) CXCL8. Here, ECM = [0.35, 0.65] ⊂ Ω = [0, 1]. Note that the initial concentrations of CXCL8, TGF-*β*, neutrophil elastase and MMPs are uniformly zero, as in experiments. *x-axis = space (the dimensionless length across the invasion chamber), y-axis = the dimensionless density/concentration of the variables.

A comparison of computational results from the mathematical model with experimental data [27] is shown in Figs 5-6. Hereafter, in order to calculate the population of cells (tumor cells, N1 TANs, N2 TANs) and level of chemical variables (CXCL-8, TGF-*β*, NET/NE, DNase, MMPs) at various times in the mathematical model, we integrate the density and concentration over the space: density of tumor cells 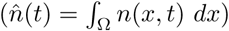, N1 TANs: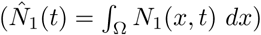, N2 TANs: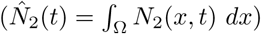, and concentrations of ECM 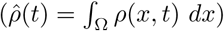, CXCL-8 (*Ĉ*(*t*) = ∫_Ω_ *C*(*x, t*) *dx*), TGF-*β*: (*Ĝ*(*t*) = ∫_Ω_ *G*(*x, t*) *dx*), NET/NE (*Ê*(*t*) = ∫_Ω_ *E*(*x, t*) *dx*), DNase I 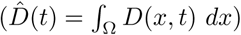, and MMPs 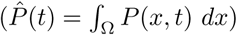. In the experiments, Park *et al*. [27] found that the presence of neutrophils in the lower chamber could enhance tumor cell invasion through NET formation and NE activities, and the DNase treatment abrogated the invasion-promoting effect of neutrophils in the lower chamber. Fig 5(A-C) shows time courses of the tumor population, population of invasive tumor cells, and neutrophil population (N1 (red solid), N2 (blue dashed) TANs), respectively, in the absence (control) and presence (^+^TAN) of neutrophils. In the presence of neutrophils, both total (Fig 5A) and invasive (Fig 5B) tumor cell populations are increased relative to the control case due to neutrophil transition (Fig 5C) and NE activities (Fig 4D) in the system. After 22*h* the number of 4T1 tumor cells invading the lower chamber almost doubled (∼190%) in the co-culture with neutrophils (red bar (^+^TAN); left panel in Fig 5D) in the lower chamber as compared to the control (blue bar; left panel in Fig 5D) in experiments [27]. In the model simulations, the number of invading tumor cells increased ∼2-fold in the presence of neutrophils in the lower chamber (red bar (^+^TAN); right panel in Fig 5D) relative to the control (absence of neutrophil (blue bar); right panel in Fig 5D). As Park *et al*. [27] note, several tumor cell lines (4T1, BT-549) invade the lower chamber even in the absence of neutrophils in the lower well, which indicates the intrinsic invasiveness of tumor cells.

**Fig 5.**
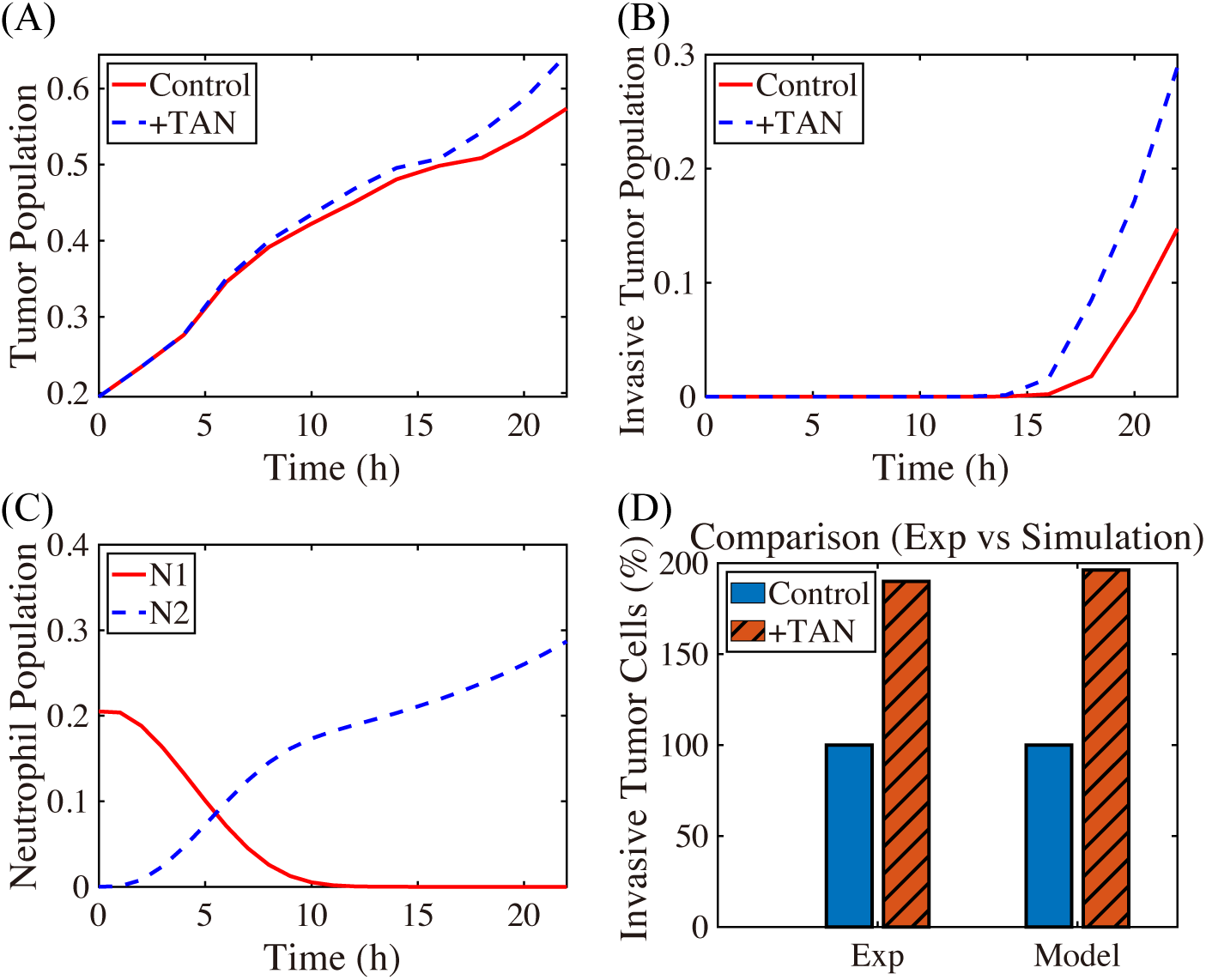
TAN-promoted cancer cell invasion (Experiment [27] & simulation). (A-B) Time courses of populations of total tumor cells (A) and invasive tumor cells (B). (C) Time courses of N1 (red solid) and N2 (blue dashed) neutrophils. (D) Experimental data from the invasion assay in [27] (left column; 4T1 cancer cells) and computational results from mathematical model (right column). The graph shows the (scaled) populations of invasive tumor cells at *t* = 22 *h* in the absence (control) or presence (^+^TAN) of neutrophils. Here and hereafter cell populations are derived from the continuum density.

In Fig. 6, we investigate the effect of DNase I against NET/NE on tumor invasion. Our mathematical model predicts that injection of DNase I in the lower chamber of the transwell can inhibit NET/NE activities (Fig. 6A) and reduce the TAN-induced invasiveness of tumor cells (right panel in Fig 6B). Park *et al*. [27] showed that NE inhibition or digestion of the DNA of the NETs by DNase I can effectively abrogated the invasion-promoting effect of TANs in the lower chamber, *i*.*e*., the number of invasive 4T1 tumor cells was reduced in the presence of the neutralizing DNase I (^+^TAN^+^D) when compared to the TAN case in the absence of the DNase I (^+^TAN) (left panel in Fig 6B). Thus, simulations are in good agreement with experimental data [27]. By definition, NETs are associated with neutrophil proteases with the extracellular histone-bound DNA [128]. In the experiments [27], pro-invasive effects of NETs were shown to be associated with protease activities of NET-associated protease, NE. Park *et al*. [27] found that the NE inhibitor reduced the extension of cancer cell-induced NETs and inhibit TAN’s ability to promote the invasion of 4T1 and BT-549 breast cancer cells. They also found that DNase I treatment can also prevent lung metastasis in mice. However, it is worth observing that this DNase I is not enough to completely inhibit the aggressive migration of tumor cells from the upper chamber to the lower chamber, since they are able to invade in the absence of neutrophils.

**Fig 6.**
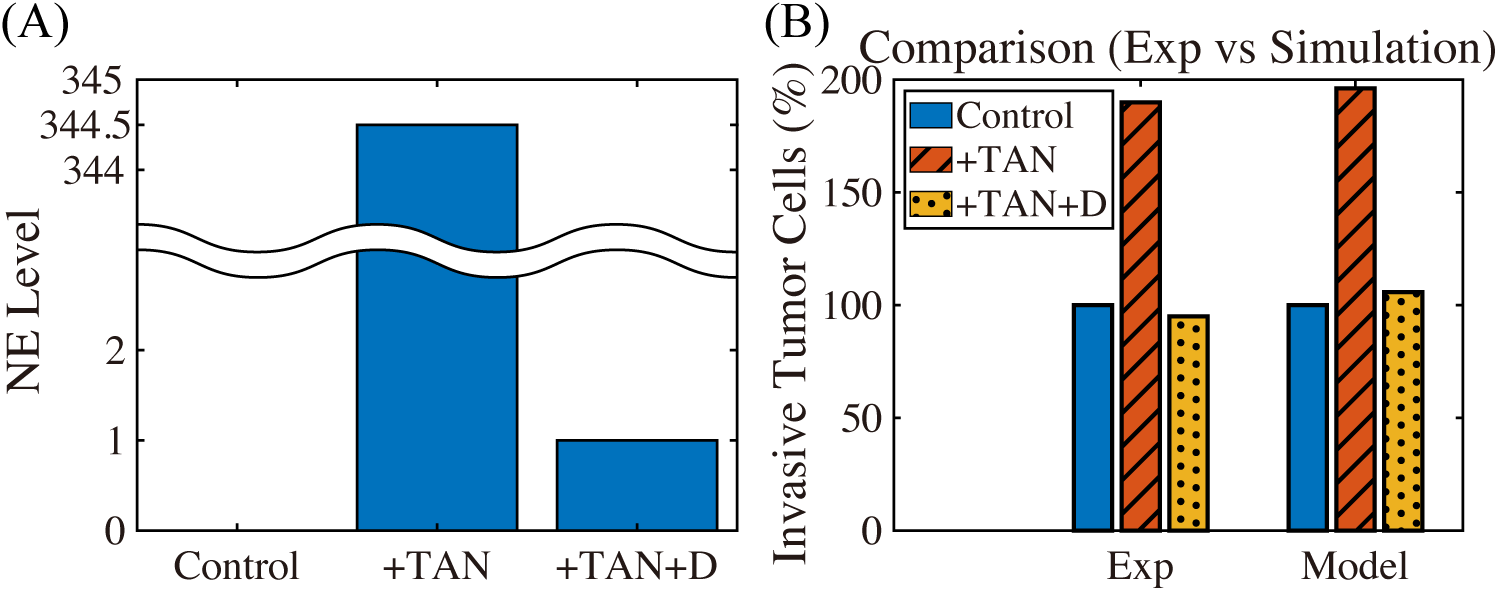
DNase I treatment against NE can abrogate the invasion-boosting effects of neutrophils (Experimental data [27] and simulation results). (A) NE levels in the system in the absence (control) and presence (^+^TAN) of neutrophils, and DNase treatment (^+^TAN^+^D) cases at *t* = 22 *h*. (B) Experimental data from the invasion assay in [27] (left column; 4T1 cancer cells) and computational results from mathematical model (right column). The graph shows the (scaled; %) population of invasive tumor cells at *t* = 22 *h* in the absence (control; blue) and presence (^+^TAN; red shaded) of neutrophils, and DNase treatment (^+^TAN^+^D; yellow dotted) cases. Addition of DNase I reduces the number of invading tumor cells by almost 50%.

The N1 → N2 transition of TANs was shown to play a critical role in promoting tumor growth, angiogenesis, invasion [3, 34, 35, 129], and ultimately metastasis initiation [130, 130, 131]. Fig 7A shows the spatial profiles of the tumor cells for various N1 → N2 transition rates (*λ*_12_ = 1.6 × 10^−4^ (red solid), 1.6 × 10^−2^ (blue dashed), 1.6 × 10^−2^ (pink with marks)) at the final time (*t* = 22 *h*). The N1- and N2-dominant spatial profiles of neutrophils in the lower chamber for the corresponding parameter set are shown in Fig 7B and Fig 7C, respectively. If we increase the rate *λ*_12_ (differentiation degree of anti-tumorigenic TANs to tumor-promoting TANs), the N2 population dominates the lower chamber (Figs 7(B-C)) with the higher population ratio N2:N1 of TANs (Fig 7F). Fig 7D shows time courses of NE levels for various values of the differentiation rate (*λ*_12_ = 1.6 × 10^−4^, 1.6 × 10^−3^, 1.6 × 10^−2^, 1.6 × 10^−1^). The corresponding populations of invasive tumor cells and neutrophils (N1,N2) at final time (*t* = 22 *h*) are shown in Fig 7E and Fig 7F, respectively. For larger *λ*_12_, more aggressive, tumor-promoting N2 TANs in the lower chamber can interact with tumor cells in the upper chamber (Fig 7A) by secreting more NE (Fig 7D). This leads to an increased tumor population and enhanced tumor cell invasion (Fig 7E). This increased invasiveness of the tumor cells is the result of the mutual interactions between tumor cells in the upper well and the neutrophils in the lower well. For instance, the NET/NE level increases as *λ*_12_ increases (Fig 7D). For large *λ*_12_, the most of N1 TANs are converted into the N2 phenotype (4th column (*λ*_12_ = 1.6 × 10^−1^) in Fig 7F), leading to efficient tumor cell migration (Fig 7E). However, when this transition rate is small (*λ*_12_ = 1.6 × 10^−4^), the less effective N1 TANs persist in the lower chamber (Fig 7B) with less population of the N2 phenotype (Fig 7C). This results in the slower (or close to zero) production of NE (Fig 7D) by TANs, and lower secretion of both TGF-*β* and MMP by tumor cells, which in turn reduces invasiveness of tumor cells by more than 48% (Fig 7E).

**Fig 7.**
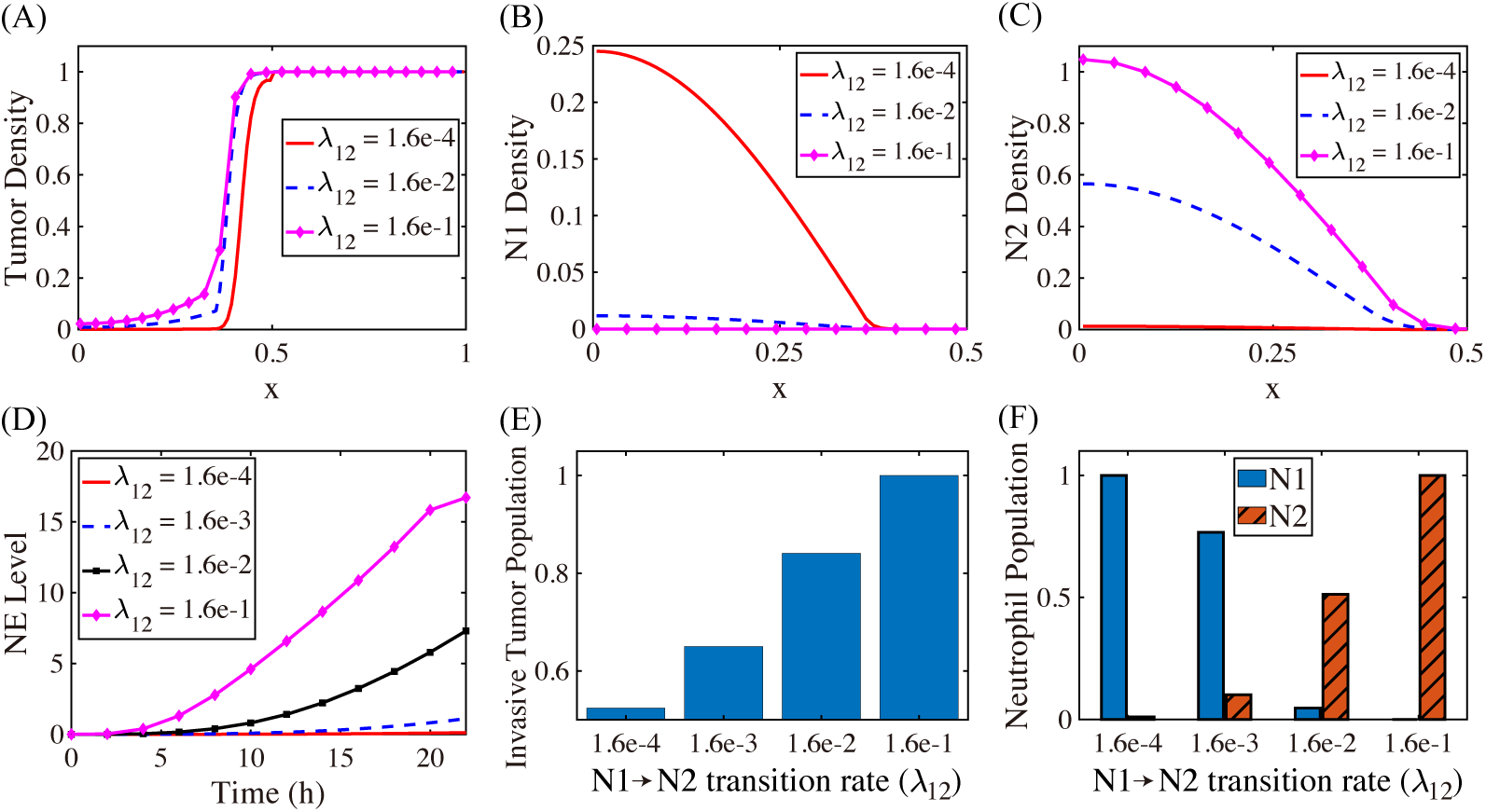
Effect of the N1 → N2 transformation on tumor invasion and N1/N2 dynamics. (A) Tumor density profiles on Ω = [0, 1] at the final time (*t* = 22 *h*) for various *λ*_12_’s (*λ*_12_ = 1.6 × 10^−4^, 1.6 × 10^−2^, 1.6 × 10^−2^). (B,C) Density profiles of the N1 and N2 TANs in the lower chamber ([0, 0.5]) for the corresponding *λ*_12_’s in (A). (D) Time courses of NE levels for various values of the differentiation rate (*λ*_12_ = 1.6 × 10^−4^, 1.6×10^−3^, 1.6 × 10^−2^, 1.6 × 10^−1^). (E,F) Scaled population of invasive tumor cells and neutrophils (N1 and N2) at the final time (*t* = 22 *h*) for various *λ*_12_’s in (D).

We study the effect of chemotaxis of tumor cells on tumor cell infiltration into the lower chamber through the ECM barrier in the middle. Fig 8 shows the tumor density (Fig 8A) and the populations of migratory tumor cells (left panel in Fig 8B) at *t* = 22 *h* as a function of the chemotactic sensitivity (*χ*_*E*_ = 4.02 × 10^−6^, 4.02 × 10^−5^, 8.02 × 10^−5^). One sees that as *χ*_*E*_ increases, the population of migratory tumor cells increases and they move faster toward the ECM membrane. We note that these tumor cells grow faster in the lower chamber due to higher level of NE and NE-mediated growth, leading to increased total tumor populations (right panel in Fig 8B).

**Fig 8.**
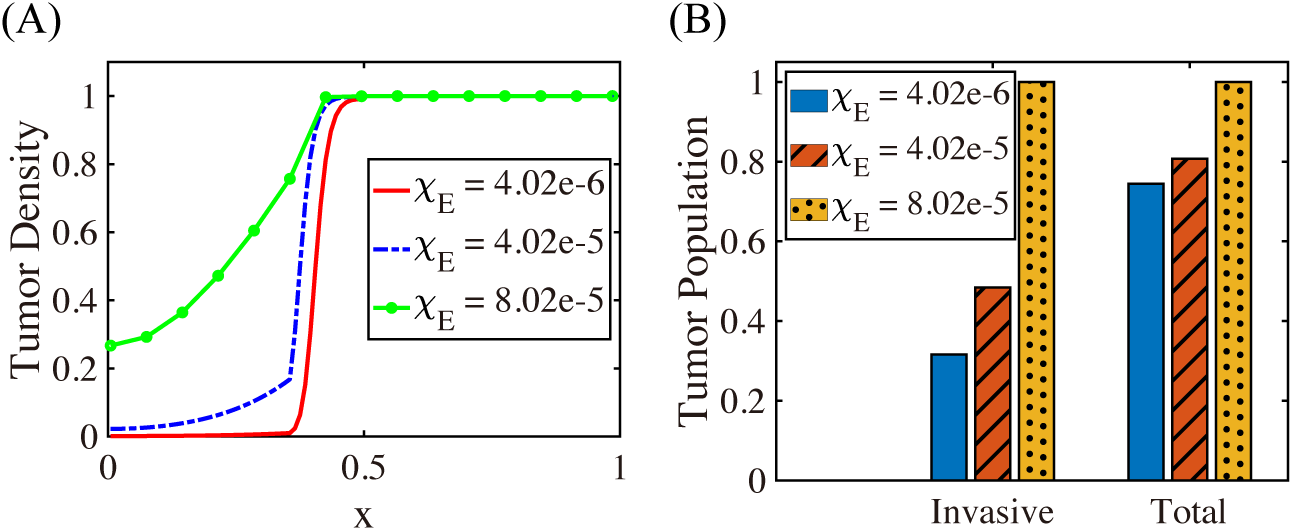
The effect of chemotaxis on tumor cell migration. (A) Spatial profiles of the tumor cell density on Ω = [0, 1] at final time (*t* = 22 *h*) for different values of the chemotactic sensitivity (*χ*_*E*_ = 4.02 10^−6^, 4.02 10^−5^, 8.02 10^−5^). (B) The (scaled) number of invasive (left panel) and total (right panel) tumor cells at final time (*t* = 22 *h*) as a function of *χ*_*E*_. As *χ*_*E*_ increases, the number of migrating tumor cells is increased.

Fig. 9A shows the spatial profiles of tumor densities at final time (*t* = 22 *h*) for various haptotactic parameters (*χ*_*ρ*_ = 1.26 × 10^−5^, 1.26 × 10^−4^, 3.26 × 10^−4^). One can observe that more tumor cells invade the lower chamber when *χ*_*ρ*_ is increased (*χ*_*ρ*_ : 1.26 × 10^−5^ → 1.26 × 10^−4^ → 3.26 × 10^−4^). The scaled numbers of migrated tumor cells at final time (*t* = 22 *h*) as a function of the haptotactic sensitivity *χ*_*ρ*_ are shown in Fig 9B. As *χ*_*ρ*_ increases, the number of migrating tumor cells and overall speed of invasion increase. We also explored the combined sensitivity of chemotactic- and haptotactic-stimulus on the invasiveness of tumor populations [56] (data not shown). We, again, found that the tumor cells show higher potential of invading the lower chamber when they are stimulated with higher chemotactic (*χ*_*E*_) and haptotactic (*χ*_*ρ*_) sensitivities. Furthermore, the combined movement at final time is more evident in the lower chamber due to the stronger degree of interaction between tumor cells and N2 TANs.

**Fig 9.**
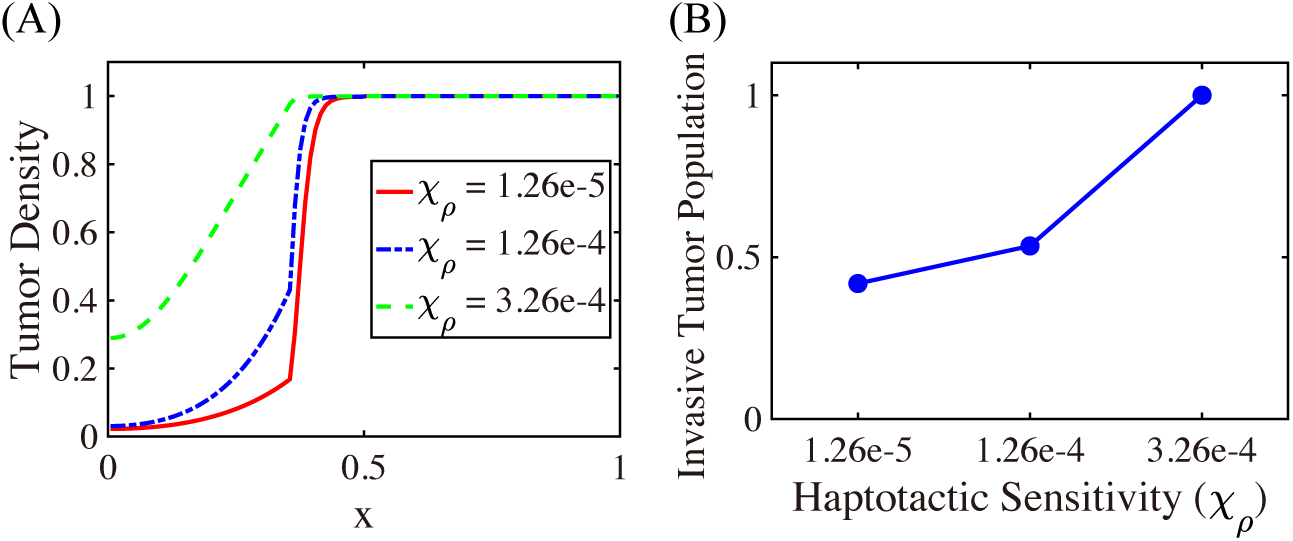
The effect of haptotaxis on invasion. (A) Spatial profiles of the tumor cell density on Ω = [0, 1] at *t* = 22 *h* for different values of the haptotactic sensitivity (*χ*_*ρ*_ = 1.26 × 10^−5^, 1.26 × 10^−4^, 3.26 × 10^−4^). (B) The population of migrating tumor cells in the lower chamber as a function of *χ*_*ρ*_ at final time (*t* = 22 *h*). As the haptotactic parameter (*χ*_*ρ*_) increases the tumor cells invade into the region occupied by the neutrophils more rapidly.

We now investigate the effect of TGF-*β* on the regulation of tumor cell invasion. TGF-*β* was shown to contribute the *N* 1 → *N* 2 conversion of TANs [47, 64], therefore, further tumor growth, and suggested to promote tumor invasion [3] and metastatic potential [29]. Fig 10 shows the TGF-*β* level and scaled population of N2 TANs and invasive tumor cells for various levels of TGF-*β* secretion rate from tumor cells (*λ*_*G*_ =0.04, 0.4, 1.0, 4.0). As *λ*_*G*_ increases, the increased TGF-*β* level (Fig 10A) promotes the *N* 1 → *N* 2 transition activities in the lower chamber, leading to the increased population of N2 TANs (Fig 10B). This results in an increase in the number of the invasive tumor cells (Fig 10C) due to increased interaction between neutrophils in the lower chamber with tumor cells in the upper chamber and faster NE-mediated chemotaxis of tumor cells.

**Fig 10.**
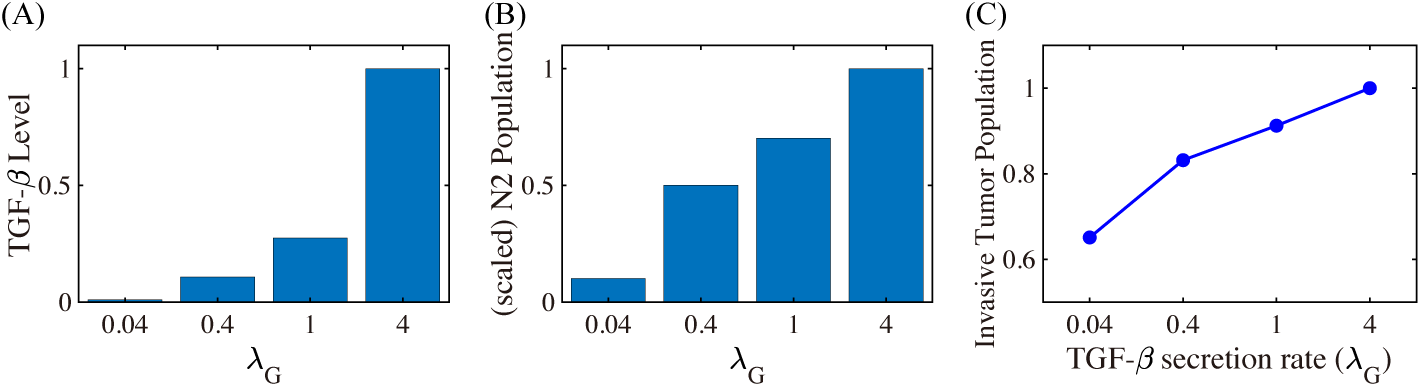
The effect of TGF-*β* on tumor cell invasion. (A) TGF-*β* level at *t* = 22 *h* for various secretion rates (*λ*_*G*_ = 0.04, 0.4, 1.0, 4.0) of TGF-*β* by tumor cells. (B,C) The scaled population of N2 TANs (B) and migrating tumor cells (C) at *t* = 22 *h* for the same set of *λ*_*G*_’s.

In Fig 11, we investigate the effect of permeability of the transfilter on the number of migrating tumor cells. In our modeling framework, the basic permeability parameters (*γ*_*c*_ for cells and larger value *γ* (*γ*_*c*_ *< γ*) for chemicals) represent the relative pore size in the membrane of the transwell for cells and chemicals [27, 31]. Two parameters (*γ*_*c*_, *γ*) were reduced or increased at the same fold. One observes that as the transfilter pore size increases (or decreases), more (or less) tumor cells in the upper chamber cross the membrane (Fig 11A), increasing (or decreasing) the overall tumor population in the lower chamber and the number of invasive tumor cells (Fig 11B). This is due to the ease of crossing the weakened physical barrier and increased proliferation rate of tumor cells via the enhanced cross-talk between tumor cells and neutrophils in the lower chamber [27].

**Fig 11.**
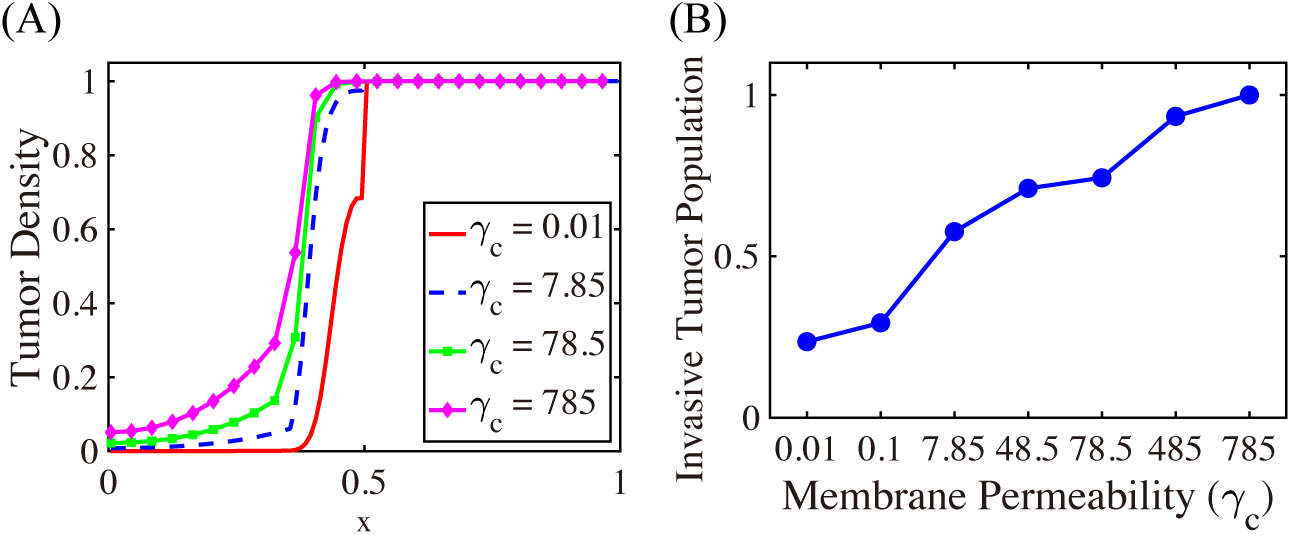
The effect of the transfilter permeability (*γ*_*c*_, *γ*) on tumor cell invasion. (A) Spatial profiles of tumor densities at the final time (*t* = 22 *h*) for various permeability parameters of the transmembrane (*γ*_*c*_ = 1.0 × 10^−2^, 7.85, 7.85 × 10^1^, 7.85 × 10^2^). (B) The (scaled) number of migratory tumor cells as a function of the permeability (*γ*_*c*_ = 1.0 × 10^−2^, 1.0 × 10^−1^, 7.85, 4.85 × 10^1^, 7.85 × 10^1^, 4.85 × 10^2^, 7.85 × 10^2^)..

### Sensitivity Analysis

The mathematical model developed in this work includes 44 parameters, some of which are available in the literature or can be reasonably estimated. However, there are some parameters for which no experimental data are known or they may affect the given system significantly. Parameters to be considered are *r, λ*_12_, *λ*_*G*_, *μ*_*ρ*1_, *μ*_*ρ*2_, *χ*_*ρ*_, *χ*_*E*_, *μ*_*ED*_, *μ*_*P M*_. In order to identify how sensitive is the cell populations and signaling levels after 12, 20 or 24 *hours* to those chosen parameters, we have performed a sensitivity analysis using a modified version of the general latin hypercube sample (LHS) and partial rank correlation coefficient (PRCC) developed in [132]. The computation was done by simulation results from the PDE model (6)-(18) with selected parameter sets and the modified Matlab files available from the website of Denise Kirschner’s Lab: http://malthus.micro.med.umich.edu/lab/usadata/. The analysis was carried out for cell populations and molecule levels from densities of all variables by integration. A physically-reasonable range ([*P*_*min*_, *P*_*max*_]) of these parameters was chosen and we divided each range into 5,000 sub-intervals of uniform length while all other parameters were fixed. For those nine parameters of interest, we calculated the PRCC value [132] in the interval [−1, 1]. The sign of the PRCC value of a parameter *P* determines whether an increase in *P* will increase (+) or decrease (-) the main variable of interest at a given time (*t* = 12, 20, 24 *h*). However, if the absolute value of the PRCC value is small, say close to zero, it doesn’t provide a significant correlation between *P* and the main variable. When the absolute value of the PRCC value is closer to 1, it statistically indicates the strong correlation between *P* and the main variable.

The computed PRCC values and their associated *p*-values of all variables for the nine perturbed parameters (*r, λ*_12_, *λ*_*G*_, *μ*_*ρ*1_, *μ*_*ρ*2_, *χ*_*ρ*_, *χ*_*E*_, *μ*_*ED*_, *μ*_*P M*_) are shown in Fig 12. Here, as before, the population of tumor cells is defined as 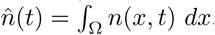, at *t* =12 (upper panel), 20 (middle panel), 24 (lower panel) hours. Populations of other cells and levels of molecules are defined in a similar fashion. Based on the PRCC results, we conclude that the tumor cell population is positively correlated to the parameters *r, μ*_*ρ*1_, *χ*_*ρ*_, but is weakly correlated with *μ*_*ρ*2_, *χ*_*E*_, *μ*_*P M*_. The tumor population is also negatively correlated to the parameter *μ*_*ED*_. While the N1 TANs is insensitive to most parameters, it is negatively correlated with the TAN transition rate (*λ*_12_) and TGF-*β* secretion rate (*λ*_*G*_). On the other hand, the N2 TAN population is positively correlated with *λ*_12_ and *λ*_*G*_, but is only weakly correlated with other parameters. One also observes that the ECM density is negatively correlated with *r, μ*_*ρ*1_, and *χ*_*ρ*_, but positively correlated with *μ*_*ED*_. One also can see that the growth rate of tumor cells (*r*) is very sensitive to the concentrations of CXCL, TGF-*β*, and ECM while *μ*_*ED*_ shows a high sensitivity for tumor cell density and concentration of ECM, CXCL, TGF-*β*, and NET/NE.

**Fig 12.**
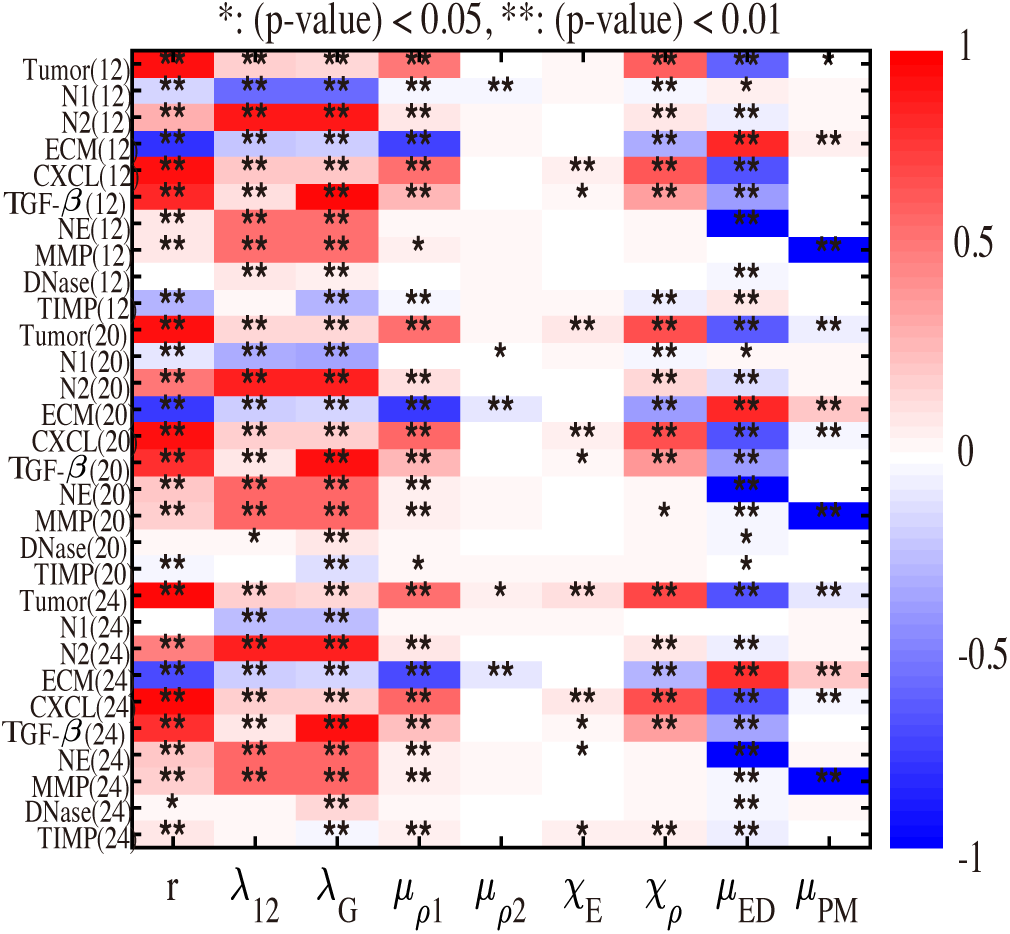
Sensitivity Analysis: General Latin Hypercube Sampling (LHS) scheme and Partial Rank Correlation Coefficient (PRCC) performed on the mathematical model. The reference output is PRCC values (red for positive PRCC values; blue for negative PRCC values) for the populations of cells (tumor cells, N1 TANs, and N2 TANs) and concentrations of molecules (tumor ECM, CXCL, TGF-*β*, NE, MMPs, DNase, and TIMP) at time *t* = 12, 20, 24 *h*.

Fig 13 shows the PRCC values of the migratory tumor cell population in the lower chamber for the same perturbed parameters (*r, λ*_12_, *λ*_*G*_, *μ*_*ρ*1_, *μ*_*ρ*2_, *χ*_*ρ*_, *χ*_*E*_, *μ*_*ED*_, *μ*_*P M*_). Here, the invasive population is defined as 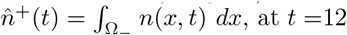, at *t* =12 (blue solid), 20 (red comb), 24 (yellow dot) hours. The numbers of invasive tumor cells are positively correlated with the parameters *r, μ*_*ρ*1_, *χ*_*ρ*_, but not so sensitive to *λ*_12_, *λ*_*G*_, *μ*_*ρ*2_, *χ*_*E*_, *μ*_*P M*_. In particular, the invasive capacity of tumor cells will increase significantly if the growth rate of tumor cells (*r*) or the ECM degradation rate (*μ*_*ρ*1_) from N2-secreted NE is increased. We note that the migration potential of tumor cell population is also positively correlated with the haptotactic strength *χ*_*ρ*_. However, the tumor cell invasiveness is negatively correlated with the degradation rate of NE by DNase I (*μ*_*ED*_). Therefore, tumor cell invasion can be sensitively reduced in response to DNase treatment.

**Fig 13.**
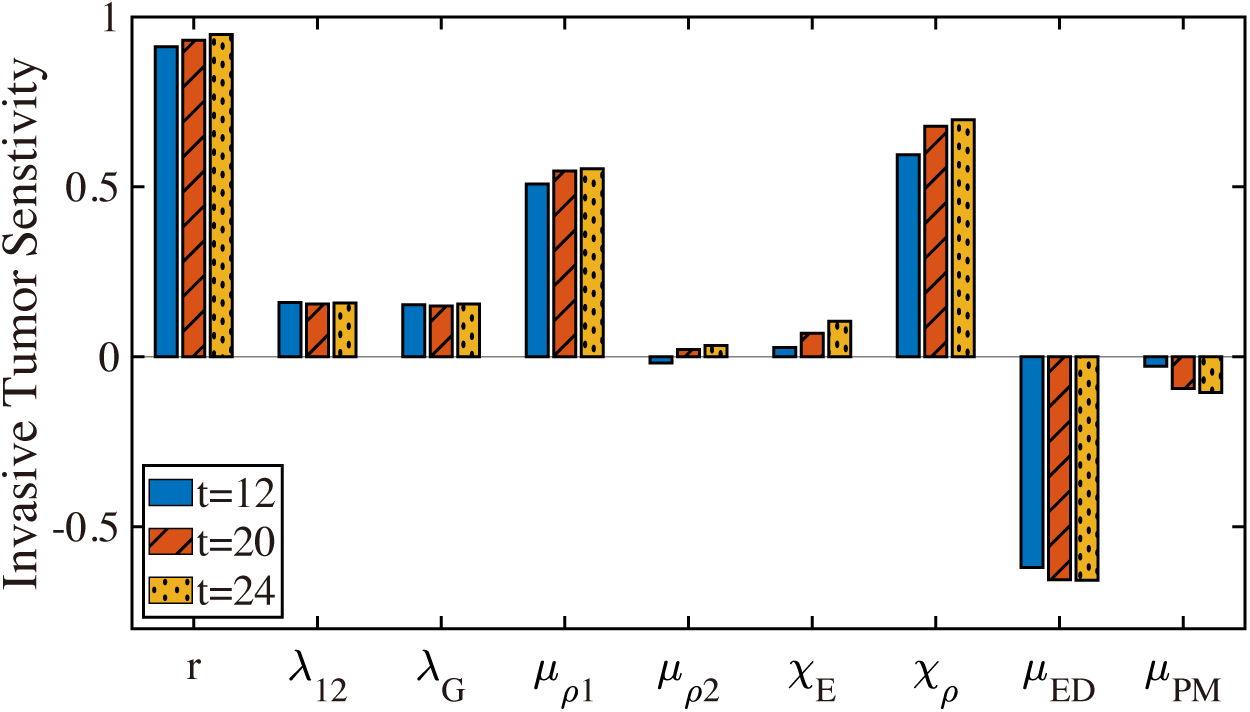
Sensitivity Analysis: PRCC values of migratory tumor cells (cell population in the lower chamber) at time *t* =12 (blue solid), 20 (red comb), 24 (yellow dot) *h*.

### Application of the model

We first test the effect of TANs on invasive and metastatic potential of tumor cells in Fig 14. To quantify the TAN-promoting effect on tumor-cell invasion, we initially placed different TAN populations on the lower chamber with various initial ratios of the TAN population to tumor population (TAN:tumor=1:100,1:10,1:2, 1:1, 2:1, 5:1, 10:1). As the ratio (TAN:tumor) increases, the invasive potential of tumor cells through the filter also increased (Fig 14A) due to the increased population of N2 TANs (Fig 14B) and increased potential of chemotactic- and haptotactic-mediated movement. For example, the up-regulation of NEs (Fig 14C) and MMPs (Fig 14D) promotes the ECM degradation (Fig 14F), increasing the haptotactic potential. On the other hand, the up-regulation of TGF-*β* (Fig 14E) enhances the critical *N* 1 → *N* 2 transition, further increasing N2 TAN activities (Fig 14B) and closely connecting the positive feedback loop between the tumor cells in the upper chamber and TANs in the lower chamber. Welch et al [45] tested the effect of Tumor-elicited PMNs (tcPMN) on tumor cell invasion by placing both TANs and tumor cells on the top of the Matrigel in the upper chamber and found that tcPMN can effectively stimulate invasive and metastatic potentials of mammary adenocarcinoma cells. For example, they found that the invasive potential of tumor cells (MTLn2) increased up to 7-fold and 25.5-fold higher for PMN:tumor ratios of 3:1 and 30:1, respectively [45]. These results suggest that the relative portion of TANs in a TME can affect the critical tumor cell invasion, therefore, metastatic potential of tumor cells, and that initial recruitment strength of TANs to the TME by tumor cells may be an important prognostic factor in determining metastatic potential in patients as suggested in experiments [10, 39, 133, 134]. In particular, neutrophil-to-lymphocyte ratio (NLR) in blood was shown to be an important prognostic factor for cancer progression [134–137] including lung cancers [9, 138].

**Fig 14.**
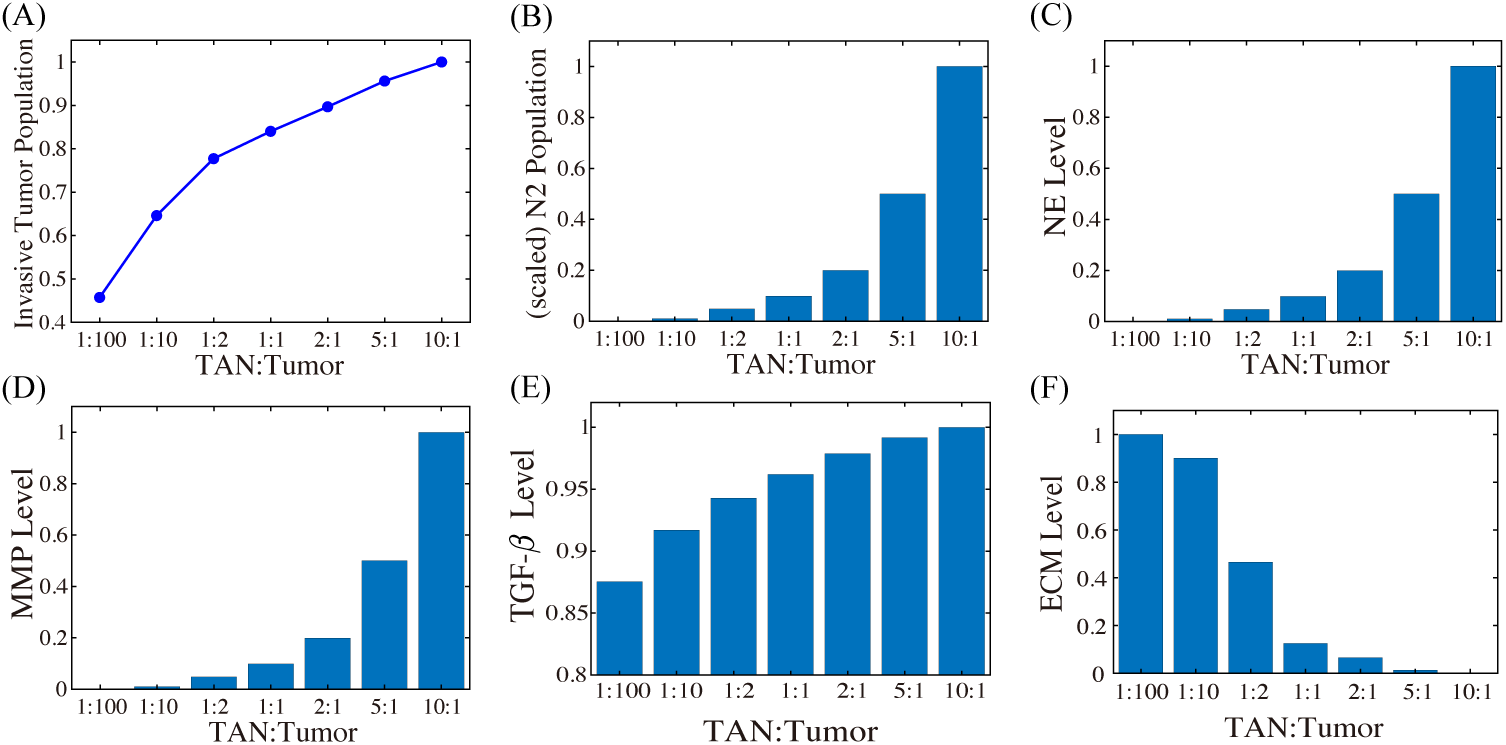
The effect of the TANs on invasive potential of tumor cells. (A) The (scaled) Invasive tumor population at final time *t* = 22 *h* with various initial ratios of the TAN population to tumor population (TAN:tumor=1:100,1:10,1:2, 1:1, 2:1, 5:1, 10:1). (B-F) Scaled N2 population (B), NE levels (C), MMP levels (D), TGF-*β* levels (E), and ECM levels on the membrane (F) corresponding to the same ratios in (A) at final time.

Blocking TGF-*β* and its receptors was shown to inhibit tumor growth [32, 139] and critical cell invasion [31, 140, 141], decrease tumourigenic potential [139, 142], and reduce metastatic incidence [143] through many different pathways [29, 144]. In order to investigate the effect of TGF-*β* suppression on tumor cell invasion. we introduce a new variable *A*(*x, t*) for the TGF-*β* anti-body and derive the following equations including the modified equation of TGF-*β* from (11)

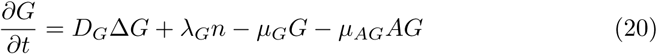

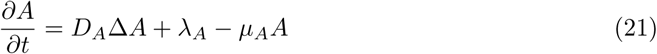

where *μ*_*AG*_ is the consumption rate of TGF-*β* due to antibody reaction, *D*_*A*_ is the diffusion coefficient of the antibody, *λ*_*A*_ is the injection rate of the antibody, and, finally, *μ*_*A*_ is the natural decay rate of the antibody. Fig 15A shows the (scaled) population of invasive tumor cells for various growth rates (*r* = 1.2, 1.3, 1.35, 1.4, 1.45, 1.5) and injection rates (*λ*_*A*_ = 0, 1.0 × 10^−2^, 1.0 × 10^−1^, 2.0 × 10^−1^, 3.0 × 10^−1^, 5.0 × 10^−1^,6.0 × 10^−1^, 7.0 × 10^−1^, 1.0, 1.0 × 10^1^) of the TGF-*β* antibody. For a fixed growth rate of tumor cells, the TGF-*β* inhibitor treatment can effectively reduce the invasiveness of tumor cells. For example, the invasive tumor population with a high dose of antibody (*λ*_*A*_ = 10) is reduced by 62% compared to the case without the antibody (*λ*_*A*_ = 0) when *r* = 1.5. However, the increasing growth rate can abrogate this antibody-induced reduction in tumor cell invasion for a fixed *λ*_*A*_ (*λ*_*A*_ = 1.2 → 1.3 → … → 1.5). DNase treatment, the NE inhibitor, was shown to reduce the NE-mediated invasion of tumor cells (4T1 and BT-549) experimentally [27] and theoretically (Fig 6). Thus, another effective therapeutic approach to slowing tumor invasion is to apply DNase I for a combination therapy (TGF-*β* inhibitor +DNase I) given its pivotal roles in tumorigenesis [4, 10, 27]. We test the efficacy of the combination therapy in Fig 15B. The invasive tumor population is reduced by 36% in response to a high dose of the TGF-*β* inhibitor (*λ*_*A*_ = 10; +Ab-D in Fig 15B) compared to control (-Ab-D in Fig 15B). Our simulation shows that blocking NE-mediated proteolytic activity of tumor cells near the membrane by DNase I in the presence of TGF-*β* inhibitor can further reduce invasiveness of tumor cells (66% reduction) invade the lower chamber (Fig 15B).

**Fig 15.**
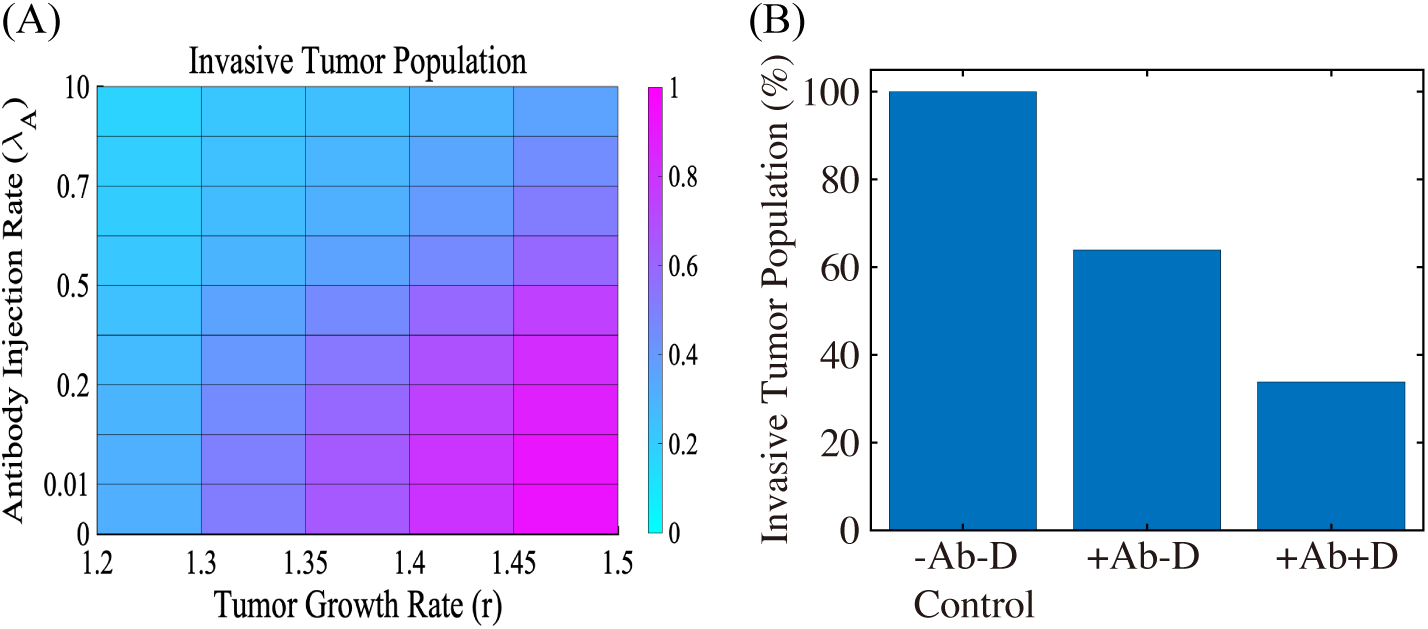
The effect of TGF-*β* blocking (+Ab) the combined therapy (+Ab+DNase I) on tumor cell invasion. (A) The (relative) population of migrating tumor cells for various growth rate (*r* ∈ [1.2, 1.5]) and injection rate (*λ*_*A*_ ∈ [0, 10]) of the TGF-*β* antibody. When TGF-*β* activity is inhibited by antibody (*r* = 1.5), fewer cells (62% reduction) invade the lower chamber. (B) Population of migrating tumor cells when TGF-*β* antibody was added in the absence (+Ab-D) and presence (+Ab+D) of DNase I relative to the control (-Ab-D). When proteolytic activity of tumor cells near the membrane is blocked by DNase I, fewer cells (66% reduction) invade the lower chamber in the presence of TGF-*β* inhibitor.

Next, we tested if MMP inhibition by TIMP can effectively reduce the TAN-mediated invasion through the transfilter. It has been shown that TIMP treatment can significantly reduce (*>* 50%) the number of migratory breast cancer cells (*>* 50%) through 8-*μm* pores in a Boyden invasion chamber assay [145]. In the mathematical model, inhibition of MMP is implemented by injecting TIMP (*λ*_*M*_ *>* 0) in Eq (15) which abrogates MMP production through a term of degradation of MMPs, 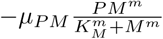, on the right hand side of Eq (14). We also tested if a combination therapy (TIMP+TGF-*β* inhibitor) can further reduce the invasion potential of tumor cells. Fig 16 shows the (relative) MMP levels, ECM levels, and number of migratory tumor cells at final time (*t* = 22 *h*) for control (-TIMP-Ab), TIMP alone (+TIMP-Ab), and combination treatment (+TIMP+Ab). One can see that TIMP treatment can inhibit the tumor cell motility by 26% (+TIMP-Ab in Fig 16C) through the significant reduction in MMP activities (82%; Fig 16A) and relatively intact ECM level (Fig 16B). An introduction of TGF-*β* antibody to the system in addition to TIMP (+TIMP+Ab)) significantly reduces the tumor cell migration through the filter (*>* 87%). Blocking tumor-TAN interactions by TGF-*β* inhibitor effectively lowered MMP levels (Fig 16A) and induced the intact ECM levels on the membrane (Fig 16B) [140, 143], contributing to this anti-invasion effect.

**Fig 16.**
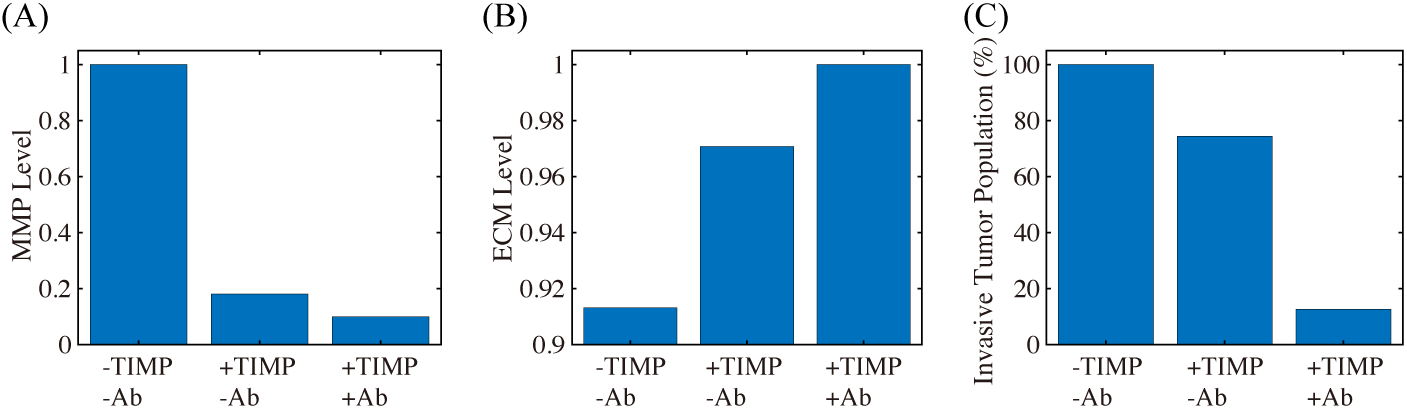
The effect of MMP blocking (+TIMP-Ab) and combined therapy (+TIMP+Ab). (A) Population of invading tumor cells when MMP activity was blocked by TIMP in the absence (+TIMP-Ab) and presence (+TIMP+Ab) of TGF-*β* antibody relative to the control (-TIMP-Ab). (B,C) Levels of MMPs and ECM corresponding to control (-TIMP-Ab), TIMP treatment (+TIMP-Ab), and combined therapy (+TIMP+Ab), respectively.

## Conclusion

TME plays an important role in regulation of tumor immunogenicity [146], tumor progression, invasion, and metastasis [3, 147]. In particular, the neutrophil-tumor interaction was shown to promote tumor cell invasion, increasing metastatic potential of cancer cells [45]. The invasion and metastasis processes of lung cancer cells may depend on many factors in TME, including immune cells and their cytokines and chemokines [50, 148]. Therefore, targeting players in TME including TANs [149] as a therapeutic approach has become more and more important [146, 150].

Experimental [27] and model simulations (Fig 5) suggest that the aggressive tumor invasion through the filter can be promoted by the mutual interaction between tumor cells in the upper chamber and TANs in the lower chamber. While the details of the *N* 1 → *N* 2 transition of TANs are still poorly understood, our model consistently suggests the pivotal role of TANs in promoting tumor cell invasion *in vitro* (Fig 7) through chemotaxis (Fig 8) and haptotaxis (Fig 9). TAN-induced signaling pathways of NET and NE were shown to actively induce tumor invasion and metastasis [42, 151, 152]. On the other hand, the presence of inhibitors of NET/NEs and MMPs was shown both experimentally to block tumor invasion [10, 27]. NET was shown to trap the circulating tumor cells (CTCs) in a lung carcinoma model, promoting tumor cell metastasis [24] by up-regulation of *β*_1_-integrin on NETs and cancer cells [153]. Thus, blocking this TAN-assisted tumor invasion can be a critical step to decrease the metastatic potential of the tumor cells. For example, DNase I treatment led to the down-regulation of NE and NET activities and reduced the invasive and metastatic potential of tumor cells (Fig 6), which is in good agreement with experimental studies [27]. Interestingly, impeding the formation of NET by DNase I treatment was shown to halt the actuation of dormant cancer cells [154].

The role of MMPs in regulation of cancer cell invasion and metastasis is well-known [55, 80]. MMP2, for instance, can not only induce tumor cell invasion by degrading ECM but promote tumor cell proliferation by enhancing vessel maturation and function in brain tumors [80]. In particular, inhibition of MMP activity can block cancer cell invasion by suppressing cell-ECM tractions and inducing cell softening [55]. In our model, TIMPs were able to partially block tumor cell invasion by heavy degradation of ECM (Fig 16A). However, when we apply combined therapeutic strategies by TIMP and TGF-*β* antibody, this effectively inhibited the tumor cell invasion through the filter (*>*87% reduction; Fig 16C). Recently, it was shown that the invasiveness of cancer cells was regulated by MMP catalytic activity via modulation of integrins-FAK signaling network [55]. Recently, TANs are shown to facilitate the metastasis to liver by increased binding activity of CCDC25 to NET DNA [155, 156]. It would be interesting to investigate the detailed mechanism of this signaling.

There are many factors that may change permeability of the narrow intercellular space for cellular infiltration. Even though the permeability parameters are fixed in *in vitro* experiments, the permeability through the narrow gap for cell invasion varies in the *in vivo* system and is regulated by cells and their cytokines/chemokines, changing the invasive potential (Fig 11). For example, TANs and NET can mediate cancer cell extravasation through TAN-CTC adhesion and breaking endothelial cell (EC) barriers, leading to active metastasis [157]. The whole process includes an initial strong adhesion process between a CTC and TANs involving selectins/integrins/ICAM1, and a series of signaling networks for CTC-EC adhesion, increased permeability from physical contraction of ECs, and the final extravasation [50, 157, 158]. A multi-scale model [159] may explain the fundamental mechanism behind this complex process in more detail by taking into account specific cell-cell adhesion [160], ECM-cell interaction [161], mechanical stress [32, 126, 141], fluid flow [162], and intracellular signaling of cellular process [32, 163].

Many studies showed that NLR in blood can be an important prognostic factor of cancer progression [134–137] including lung cancers [9, 138]. Our simulation results suggest that the presence of a larger portion of TANs in a TME (or high NLR in blood) can effectively stimulate tumor cell invasion and increase the metastatic potential through increased mutual interaction between tumor cells and N2 TANs (Fig 14). TGF-*β* mediates this critical *N* 1 → *N* 2 transition of TANs and promotes tumor growth and invasion through the filter (Fig 10). Therefore, the early detection of initial recruitment of TANs to the TME, for example by calculating NLR, may be an important step in decreasing metastatic potential in patients [10, 39, 133, 134]. Further studies on specific downstream pathways of this CCDC25-ILK-*β*-Parvin signalling [156] would be needed for development of anti-metastatic drugs that target and block the NET-cancer interaction.

Blocking TGF-*β* and its receptors was suggested to a therapeutic approach due to their ability to inhibit tumor growth [32, 139] and critical cell invasion [31, 140, 141], decrease tumourigenic potential [139, 142], and reduce metastatic dissemination [143] through various different pathways including the SMADs family [29, 144]. We showed that inhibition of TGF-*β* can effectively decrease migration potential of tumor cell cells through the transfilter by reducing the critical interaction between neutrophils and tumor cells (Fig 10, Fig 15). This illustrates the critical role of neutrophils in tumor cell invasion and importance of TGF-*β* inhibition in suppressing cell infiltration and metastasis potential. A combined approach with TGF-*β* inhibitor and DNase I can further reduce the invasiveness of tumor cells through the filter (Fig 15). We note, however, that TGF-*β* treatment, not TGF-*β* inhibition, can enhance anti-tumor efficacy through temporal immune suppression in other approaches. For example, Han *et al*. [164] showed that pretreatment with TGF-*β* prior to oncolytic virus (OV) therapy effectively inhibited tumor growth by suppressing resident microglia and natural killer (NK) cells in glioblastoma therapy trial. In the same vein, Kim *et al*. [165] also experimentally and theoretically showed that physical deletion of resident NK cells (^−^*NK*) in the TME, unexpectedly, induced better anti-tumor efficacy relative to the control case in the OV-bortezomib combination therapy since residential NK cells, if not removed, also killed infected tumor cells and depletion of NK cells significantly increased OV-mediated tumor killing. It would be interesting to see how this TGF-*β*-mediated immune suppression affects the tumor invasion and metastasis processes in these combination therapies.

We plan to investigate the role of the continuous spectrum of the *N* 1 → *N* 2 transition and the role of TANs in the regulation of NET-mediated metastasis in future work. Signaling between cells is an integral process in controlling invasive and metastatic potential of tumor cells due to unexpected mutations and chromosomal changes. This signaling often involves indirect communications between various immune cells and spatially-separated tumor cells in the TME. For example, the detailed communication between a tumor at a distant site and neutrophils in bone marrow is poorly understood. Intra-tumor heterogeneity from packing density of various cells in TME and large anisotropic transport through the tissue can affect the signaling pathways [166], thus cancer treatment [167], but despite its importance, corresponding experimental data on signaling are insufficient. Interestingly, the aging TME was recently suggested to influence tumor progression including the critical tumor cell invasion [71]. Thus, *in silico* studies on the effects of these critical interactions on cancer cell invasion, and on the responsiveness of the predictions to physical parameters, can shed insights into guiding experiments aimed at the development of new therapeutic strategies.

## Supporting information

**S1 Appendix. Parameter estimation and nondimensionalization**.

## Acknowledgments

This paper is supported by the Basic Science Research Program through National Research Foundation of Korea (NRF) funded by Ministry of Science, ICT and Future Planning: (2018R1A2B6007288) (Y.K.).

